# *Leishmania naiffi* and *Leishmania guyanensis* reference genomes highlight genome structure and gene evolution in the *Viannia* subgenus

**DOI:** 10.1101/233148

**Authors:** Simone Coughlan, Ali Shirley Taylor, Eoghan Feane, Mandy Sanders, Gabriele Schonian, James A. Cotton, Tim Downing

## Abstract

The unicellular protozoan parasite *Leishmania* causes the neglected tropical disease leishmaniasis, affecting 12 million people in 98 countries. In South America where the *Viannia* subgenus predominates, so far only *L. (Viannia) braziliensis* and *L. (V*.*) panamensis* have been sequenced, assembled and annotated as reference genomes. Addressing this deficit in molecular information can inform species typing, epidemiological monitoring and clinical treatment. Here, *L. (V*.*) naiffi* and *L. (V*.*) guyanensis* genomic DNA was sequenced to assemble these two genomes as draft references from short sequence reads. The methods used were tested using short sequence reads for *L. braziliensis* M2904 against its published reference as a comparison. This assembly and annotation pipeline identified 70 additional genes not annotated on the original M2904 reference. Phylogenetic and evolutionary comparisons of *L. guyanensis* and *L. naiffi* with ten other *Viannia* genomes revealed four traits common to all *Viannia*: aneuploidy, 22 orthologous groups of genes absent in other *Leishmania* subgenera, elevated TATE transposon copies, and a high NADH-dependent fumarate reductase gene copy number. Within the *Viannia*, there were limited structural changes in genome architecture specific to individual species: a 45 Kb amplification on chromosome 34 was present in all bar *L. lainsoni, L. naiffi* had a higher copy number of the virulence factor leishmanolysin, and laboratory isolate *L. shawi* M8408 had a possible minichromosome derived from the 3’ end of chromosome 34. This combination of genome assembly, phylogenetics and comparative analysis across an extended panel of diverse *Viannia* has uncovered new insights into the origin and evolution of this subgenus and can help improve diagnostics for leishmaniasis surveillance.

## Introduction

Most cutaneous (CL) and mucocutaneous leishmaniasis (MCL) cases in the Americas are the result of infection by *Leishmania* parasites belonging to the *Viannia* subgenus. The complexity of the molecular, epidemiological and ecological challenges associated with *Leishmania* in South America remains opaque due to our limited understanding of the biology of *Viannia* parasites. Nine *Viannia* (sub)species have been described so far: *L*. (*V*.) *braziliensis, L. (V*.*) peruviana, L*. (*V*.) *guyanensis, L*. (*V*.) *panamensis, L*. (*V*.) *shawi, L*. (*V*.) *lainsoni, L*. (*V*.) *naiffi, L*. (*V*.) *lindenbergi* and *L*. (*V*.) *utingensis*. CL and MCL are endemic in 18 out 20 countries in the Americas [1] and are mainly associated with *L. braziliensis, L. guyanensis*, and *L. panamensis*, whose frequency varies geographically. Other species are less frequently associated with human disease, and some are restricted to certain areas [2].

Human CL is partially driven by transmission from sylvatic and peridomestic mammalian reservoirs [3], via sand flies of the genus *Lutzomyia* (*sensu* Young and Duncan, 1994) in the Americas, distinct from *Phlebotomus* sand flies in the Old World [4]. Although CL has spread to domestic and peridomestic niches due to migration, new settlements and deforestation [5-7], there is still a high incidence of some *Leishmania* in sylvatic environments, such that human infection is accidentally acquired due to sand fly bites when handling livestock [8]. *L. naiffi* and *L. guyanensis* are among the *Viannia* species that show variable responses to treatment, and diversity in the types of clinical manifestations presented, and are adapting to environmental niche and transmission changes driven by humans.

*L. naiffi* was formally described from a parasite isolated in 1989 from its primary reservoir, the nine-banded armadillo (*Dasypus novemcinctus*), in Pará state of northern Brazil [9-11]. *L. naiffi* was initially placed in the *Viannia* subgenus based on its molecular and immunological characteristics [9]. Many phlebotomine species are likely to participate in the transmission of *L. naiffi* in Amazonia [12], including *Lu. (Psathyromyia) ayrozai* and *Lu. (Psychodopygus) paraensis* in Brazil [13], *Lu. (Psathyromyia) squamiventris* and *Lu. tortura* in Ecuador [14], and *Lu. trapidoi* and *Lu. gomezi* in Panama [30]. *L. naiffi* has been isolated from humans and armadillos [9-10], and detected in *Thrichomys pachyurus* rodents found in the same habitat as *D. novemcinctus* in Brazil [16]. The nine-banded armadillo is hunted, handled and consumed in the Americas and is regarded as a pest [11,17-18]. People in the same vector range as these armadillos could be exposed to infective sand flies: three *L. naiffi* CL cases followed contact with armadillos in Suriname [19]. *L. naiffi* causes localised CL in humans with small discrete lesions on the hands, arms or legs [10,20-21], which has been observed in Brazil, French Guiana, Ecuador, Peru and Suriname [19,22]. CL due to *L. naiffi* usually responds to treatment [10,22] and can be self-limiting [23], though poor response to antimonial or pentamidine therapy was reported in two patients in Manaus, Brazil [20].

*L. guyanensis* was first described in 1954 [24] and its primary hosts are the forest dwelling two-toed sloth (*Choloepus didactylus*) and the lesser anteater *Tamandua tetradactyl* [25]. Potential secondary reservoirs of *L. guyanensis* are *Didelphis marsupialis* (the common opossum) [26,27], rodents from the genus *Proechimys* [25], *Marmosops incanus* (the grey slender opposum) [28] in Brazil, and *D. novemcinctus* [29]. *Lu. umbratilis, Lu. anduzei* and *Lu. whitmani* are prevalent in forests [30] and act as vectors of *L. guyanensis* [31-33]. *L. guyanensis* has been found in French Guiana, Bolivia, Brazil, Colombia, Guyana, Venezuela, Ecuador, Peru, Argentina and Suriname [34-39].

More precise genetic screening of *Viannia* isolates is necessary to trace hybridisation between species. Infection of humans, dogs and *Lu. ovallesi* with *L. guyanensis/L. braziliensis* hybrids was reported in Venezuela [40-41]. A *L. shawi/L. guyanensis* hybrid causing CL was detected in Amazonian Brazil [42], and *L. naiffi* has produced viable progeny with *L. lainsoni* [43] and *L. braziliensis* (Elisa Cupolillo, unpublished data). There is extensive evidence of interbreeding among *L. braziliensis* complex isolates, including more virulent *L. braziliensis/L. peruviana* hybrids with higher survival rates within hosts *in vitro* [44].

*Leishmania* genomes are characterised by several key features. Genes are organised as polycistronic transcription units that have a high degree of synteny across *Leishmania* species [45]. These polycistronic transcription units are co-transcribed by RNA polymerase II as polycistronic pre-mRNAs that are 5’-transpliced and 3’-polyadenylated [46,47]. This means translation and stability of these mature mRNAs determines gene expression rather than transcription rates. In addition, *Leishmania* display extensive aneuploidy, frequently possess extrachromosomal amplifications driven by homologous recombination at repetitive sequences, and have variable gene copy numbers [48]. The *Leishmania* subgenus genomes of *L. infantum, L. donovani*, and *L. major* have 36 chromosomes [49], whereas *Viannia* genomes have 35 chromosomes due to a fusion of chromosomes 20 and 34 [45,50]. In contrast to the species of the *Leishmania* subgenus, *Viannia* parasites possess genes encoding functioning RNA interference (RNAi) machinery that may mediate infective viruses and transposable elements [51].

Fully annotated genomes have been described in detail for only two *Viannia* species: *L. panamensis* [51] and *L. braziliensis* [45,48], limiting our comprehension of their evolutionary origin, genetic diversity and functional adaptations. Consequently, we present reference genomes for *L. guyanensis* LgCL085 and *L. naiffi* LnCL223 to address these critical gaps. These new annotated reference genomes were compared to other *Viannia* species genomes to examine structural variation, sequence divergence, gene synteny and chromosome copy number changes. We contrasted the genomic configuration of *L. guyanensis* LgCL085 and *L. naiffi* LnCL223 with the *L. braziliensis* MHOM/BR/1975/M2903 assembly, two unannotated *L. peruviana* chromosome-level scaffold assemblies [52], the *L. panamensis* MHOM/PA/1994/PSC-1 reference and the *L. braziliensis* MHOM/BR/1975/M2904 reference. Furthermore, we assessed aneuploidy in five unassembled *Viannia* datasets originally isolated from humans, armadillos and primates, which are commonly used in studies on *Viannia* parasites [53–56]: *L. shawi* reference isolate MCEB/BR/1984/M8408 also known as IOC_L1545, *L. guyanensis* MHOM/BR/1975/M4147 (iz34), *L. naiffi* MDAS/BR/1979/M5533 (IOC_L1365), *L. lainsoni* MHOM/BR/1981/M6426 (IOC_L1023), *L. panamensis* MHOM/PA/1974/WR120 [53] (IOC stands for Instituto Oswaldo Cruz).

## Results

### Genome assembly from short-reads

The genomes of *L. (Viannia) guyanensis* LgCL085 and *L. (V*.*) naiffi* LnCL223 were assembled from short reads, along with an assembly of *L. braziliensis* M2904 generated in the same way as a positive control [48] (Table 1). This facilitated comparison with the published M2904 genome, which was assembled by capillary sequencing of a plasmid clone library together with extensive finishing work and with fosmid end sequencing [45], so that the ability of short reads to correctly and comprehensively resolve *Leishmania* genome architecture could be quantified.

**Table 1:**
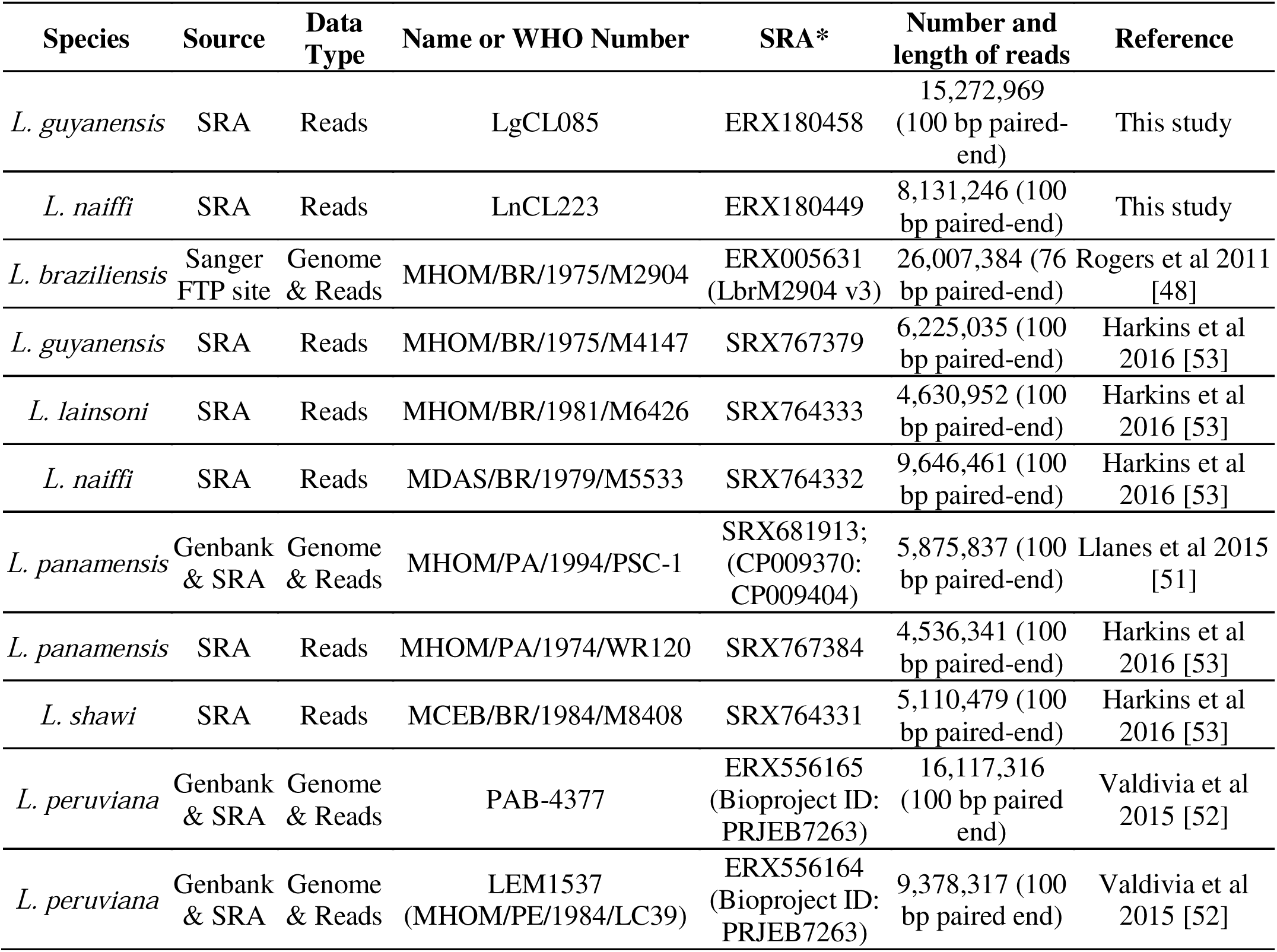
Data used in this study. The World Health Organisation (WHO) numbers are structured such that M is mammal, R is reptile, HOM is *Homo*, CAN is canine, DAS is *Dasypus* (an armadillo), CEB is *Cebus* (a primate), ARV is *Arvicanthis* (a rodent), TAR is *Tarentolae* and LAT is *Latastia* (a long-tailed lizard). The top two rows indicate the isolates for *L. guyanensis* and *L. naiffi* genomes published here. * SRA stands for SRA or TriTrypDB accession ID.

Firstly, the *L. guyanensis* LgCL085, *L. naiffi* LnCL223 and the *L. braziliensis* M2904 control reads were filtered to remove putative contaminant sequences identified by aberrant GC content, trimmed at the 3’ ends to remove low quality bases, and PCR primer sequences were removed (see Methods for details) resulting in 26,067,692 properly paired reads for *L. guyanensis*, 13,979,628 for *L. naiffi*, 34,592,618 for the *L. (V*.*) braziliensis* control (Table S1). These filtered reads for *L. guyanensis, L. naiffi* and *L. braziliensis* were *de novo* assembled into contigs using Velvet [57] with k-mers of 61 for *L. guyanensis*, 43 for *L. naiffi* and 43 for the *L. braziliensis* control optimised for each library.

The initial contigs were scaffolded using read pair information with SSPACE [58] to yield 2,800 *L. guyanensis* scaffolds with an N50 of 95.4 Kb, 6,530 *L. naiffi* scaffolds with an N50 of 24.3 Kb, and 3,782 *L. braziliensis* scaffolds with an N50 of 20.6 Kb (Table 2). The corrected scaffolds for *L. guyanensis, L. naiffi* and the *L. braziliensis* control were contiguated (aligned, ordered and oriented) using the extensively finished *L. braziliensis* M2904 reference with ABACAS [59]. The output was split into 35 pseudo-chromosomes and REAPR [60] broke scaffolds at possible misassemblies to assess contiguation accuracy. The pseudo-chromosome lengths of each sample approximated the length of each corresponding *L. braziliensis* M2904 reference chromosome with the exceptions of shorter *L. guyanensis* chromosomes 2, 4, 12 and 21, and a longer *L. naiffi* chromosome 1 (Figure S1). Post-assembly alignment of all bin contigs using BLASTn identified 44 *L. guyanensis* sequences spanning 4,566,791 bp as putative contaminants that were removed: half had high similarity to bacterium *Niastella koreensis* (Table S2).

**Table 2:**
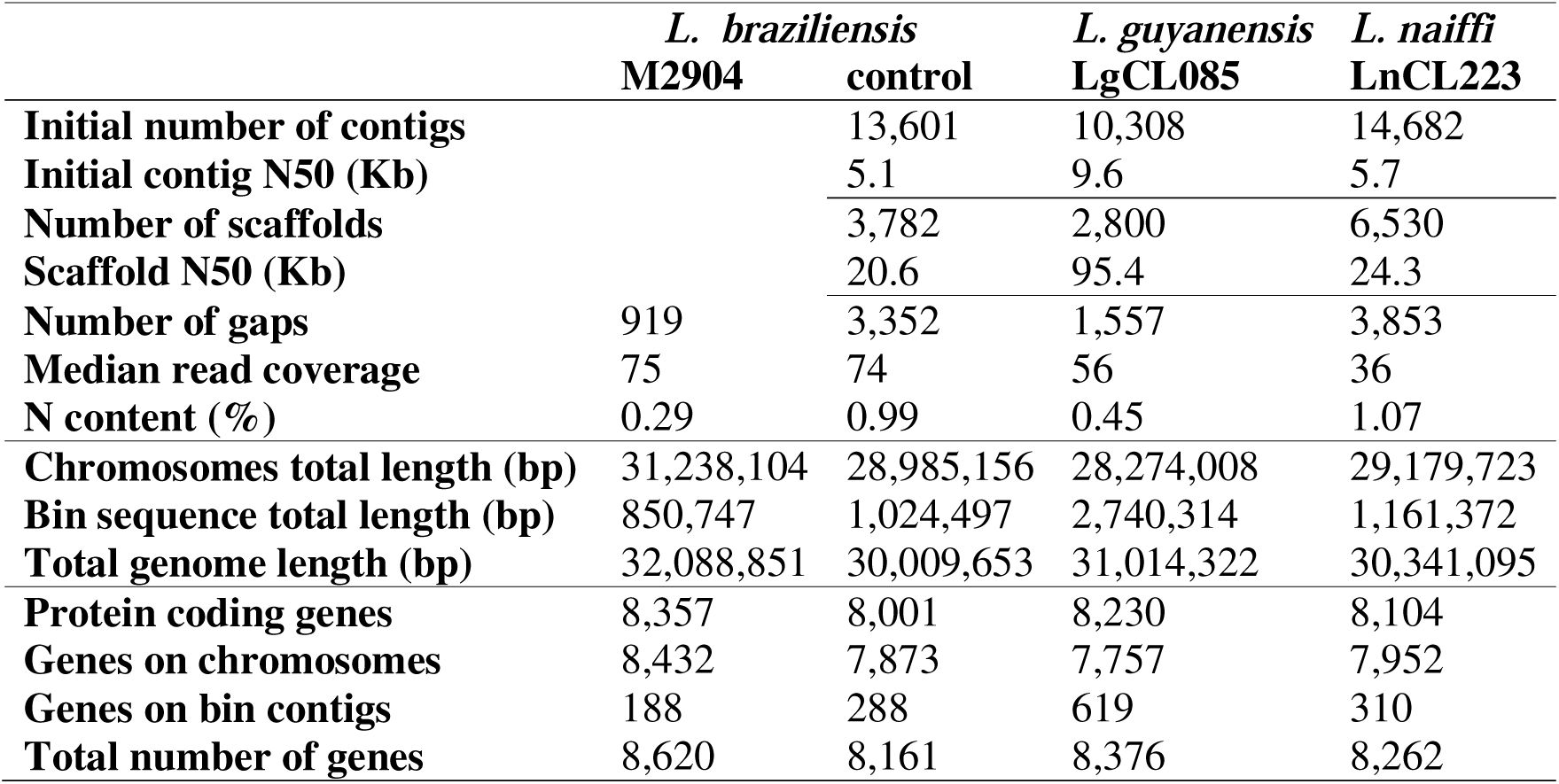
Summary of *L. braziliensis* reference M2904, *L. braziliensis* control, *L. guyanensis* LgCL085, and *L. naiffi* LnCL223 genome assembly contigs, scaffolds, gaps, read coverage, assembled chromosomal and contig sequence, and levels of gene annotation.

|  | *L. braziliensis*<br>M2904 | *L. braziliensis*<br>control | *L. guyanensis*<br>LgCL085 | *L. naiffi*<br>LnCL223 |
| --- | --- | --- | --- | --- |
| **Initial number of contigs** |  | 13,601 | 10,308 | 14,682 |
| **Initial contig N50 (Kb)** |  | 5.1 | 9.6 | 5.7 |
| **Number of scaffolds** |  | 3,782 | 2,800 | 6,530 |
| **Scaffold N50 (Kb)** |  | 20.6 | 95.4 | 24.3 |
| **Number of gaps** | 919 | 3,352 | 1,557 | 3,853 |
| **Median read coverage** | 75 | 74 | 56 | 36 |
| **N content (%)** | 0.29 | 0.99 | 0.45 | 1.07 |
| **Chromosomes total length (bp)** | 31,238,104 | 28,985,156 | 28,274,008 | 29,179,723 |
| **Bin sequence total length (bp)** | 850,747 | 1,024,497 | 2,740,314 | 1,161,372 |
| **Total genome length (bp)** | 32,088,851 | 30,009,653 | 31,014,322 | 30,341,095 |
| **Protein coding genes** | 8,357 | 8,001 | 8,230 | 8,104 |
| **Genes on chromosomes** | 8,432 | 7,873 | 7,757 | 7,952 |
| **Genes on bin contigs** | 188 | 288 | 619 | 310 |
| **Total number of genes** | 8,620 | 8,161 | 8,376 | 8,262 |

When the reads for each were mapped to its own assembled genome, the median read coverage was 56 for *L. guyanensis*, 36 for *L. naiffi* and 75 for the *L. braziliensis* control. The latter was on par with the 74-fold median coverage observed when M2904 short reads were mapped to the *L. braziliensis* reference [45,48] (Table S3). The differing coverage levels correlated with the numbers of gaps in the final genome assembly of *L. guyanensis* (1,557, Table 2) and *L. naiffi* (3,853).

### MLSA of *L. guyanensis* LgCL085 and *L. naiffi* LnCL223 with the *Viannia* subgenus

As a first step in investigating the genetic origins of these isolates, we examined their species identity using MLSA (multi-locus sequencing analysis). Four housekeeping gene sequences published for 95 *Viannia* isolates including *L. braziliensis, L. lainsoni, L. lindenbergi, L. utingensis, L. guyanensis, L. shawi* and *L. naiffi* [56] were compared with orthologs of each gene extracted from assemblies of *L. naiffi* LnCL223, *L. guyanensis* LgCL085, the *L. braziliensis* reference, *L. panamensis* PSC-1 and *L. peruviana* PAB-4377. Among the 95 were four samples with reads available [53]: *L. shawi* MCEB/BR/1984/M8408 (IOC_L1545), *L. guyanensis* MHOM/BR/1975/M4147 (iz34), *L. naiffi* MDAS/BR/1979/M5533 (IOC_L1365) and *L. lainsoni* MHOM/BR/1981/M6426 (IOC_L1023). The genes were aligned using Clustal Omega v1.1 [61] to create a network for the 102 isolates with SplitsTree v4.13.1 [62]. This replicated the expected highly reticulated structure [56] where *L. braziliensis* M2904 and *L. peruviana* PAB-4377 were in the *L. braziliensis* cluster (Figure 1).

**Figure 1:**
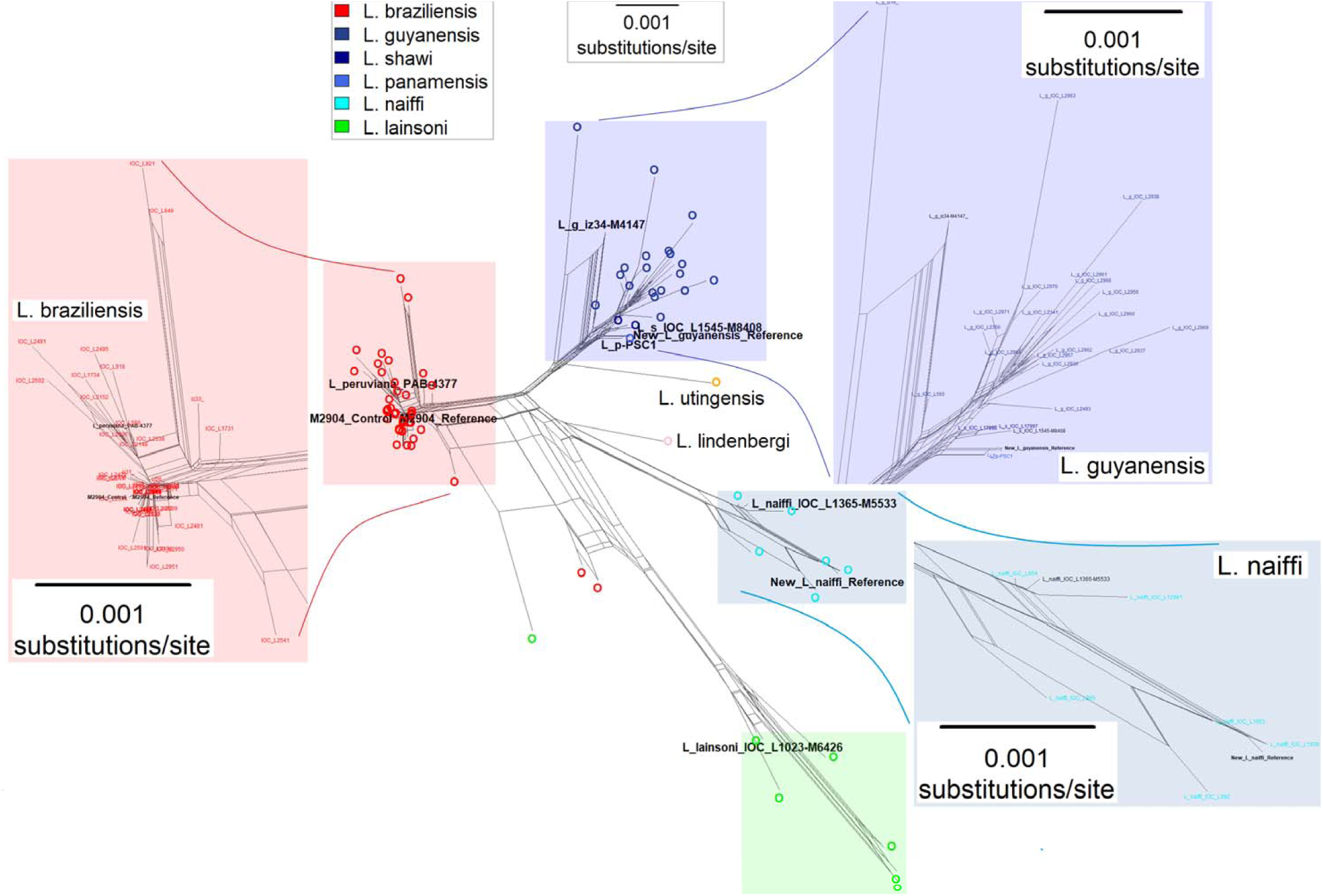
Middle: A neighbor-Net network of the uncorrected p-distances from concatenated 2,902-base sequences from four housekeeping genes for 102 *Viannia* samples. The genes were glucose-6-phosphate dehydrogenase (G6PD), 6-phosphogluconate dehydrogenase (6PGD), mannose phosphate isomerase (MPI) and isocitrate dehydrogenase (ICD). *L. naiffi* LnCL223 (cyan) is “New_L_naiffi_Reference” and is related to M5533 (IOC_L1365). *L. guyanensis* LgCL085 (blue) is “New_L_guyanensis_Reference” and is related to the *L. shawi* M8408 (IOC_L1545) assembly and the *L. panamensis* PSC-1 genome, but less so to *L. guyanensis* M4147 (iz34). The *L. braziliensis* M2904 reference and control are “M2904_Reference” and “M2904_Control”, proximal to *L. peruviana* PAB-4377. *L. lainsoni* M6426 (IOC_L1023) (green), *L. utingensis* (orange) and *L. lindenbergi* (pink) are shown. The isolate names and detail for each species complex is shown by insets in red (*L*. braziliensis), dark blue (*L. guyanensis*) and light blue (*L. naiffi*). For detailed viewing, the nexus file can be downloaded at https://figshare.com/s/eecf1c6b42ac4deb6acc and high-resolution PDF at https://doi.org/10.6084/m9.figshare.5687329.

Previous work suggests that the *L. guyanensis* species complex includes *L. panamensis* and *L. shawi* because they show little genetic differentiation from one another [56,63-65]. The MLSA here showed that the new *L. guyanensis* LgCL085 reference clustered phylogenetically in the *L. guyanensis* species complex, had no sequence differences compared to *L. panamensis* PSC-1, and seven differences versus *L. shawi* M8408 across the 2,902 sites aligned (Figure 1). *L. guyanensis* LgCL085 grouped with isolates classified as zymodeme Z26 by multilocus enzyme electrophoresis (MLEE) associated with *L. shawi* [54]. This was supported by the number and the alleles of genome-wide SNPs called using reads mapped to the *L. braziliensis* M2904 reference for *L. guyanensis* (355,267 SNPs), *L. guyanensis* M4147 (326,491), *L. panamensis* WR120 (294,459) and *L. shawi* M8408 (296,095) (Table S4).

The *L. naiffi* LnCL223 was closest to *L. naiffi* ISQU/BR/1994/IM3936, with two differences. It clustered with MLEE zymodeme Z49 based on the correspondence between the MLSA network and previously typed zymodemes, though *L. naiffi* is associated with more zymodemes than other *Viannia*. The number and the alleles of genome-wide SNPs called using reads mapped to the *L. braziliensis* reference were similar for *L. naiffi* (548,256) and M5533 (633,560) (Table S4) and consistent with the MLSA genetic distances.

There was no evidence of recent gene flow between these three species at any genome-wide 10 Kb segment and *L. naiffi* LnCL223 had fewer SNPs compared to *L. braziliensis* M2904 than *L. guyanensis* LgCL085 (Figure S2). Linking the MLSA network topology with previous work [56,63-65], four genetically distinct species complexes are represented by the genome-sequenced *Viannia* at present: (i) *braziliensis* including *L. peruviana*, (ii) *guyanensis* including *L. panamensis* and *L. shawi*, (iii) *naiffi*, and (iv) *lainsoni* (Table S4), and the less explored (v) *lindenbergi* and (vi) *utingensis* complexes (Figure 1).

### Ancestral diploidy and constitutive aneuploidy in *Viannia*

The normalised chromosomal coverage of the *L. guyanensis* LgCL085 and *L. naiffi* LnCL223 reads mapped to *L. braziliensis* M2904 showed aneuploidy on a background of a diploid nuclear genome (Figure 2). The coverage levels of reads for *L. peruviana* LEM1537, *L. peruviana* PAB-4377, *L. panamensis* PSC-1 and the triploid *L. braziliensis* control mapped to the M2904 reference, confirmed previous work (Figure S3), including the *L. braziliensis* control (Figure S4), and demonstrated that assemblies from short read data were sufficient to estimate chromosome copy number differences. Repeating this for *L. shawi* M8408, *L. naiffi* M5533, *L. guyanensis* M4147, *L. panamensis* WR120 and *L. lainsoni* M6426 showed that all these *Viannia* were predominantly disomic and thus diploidy was the likely ancestral state of this subgenus (Figure 2).

**Figure 2:**
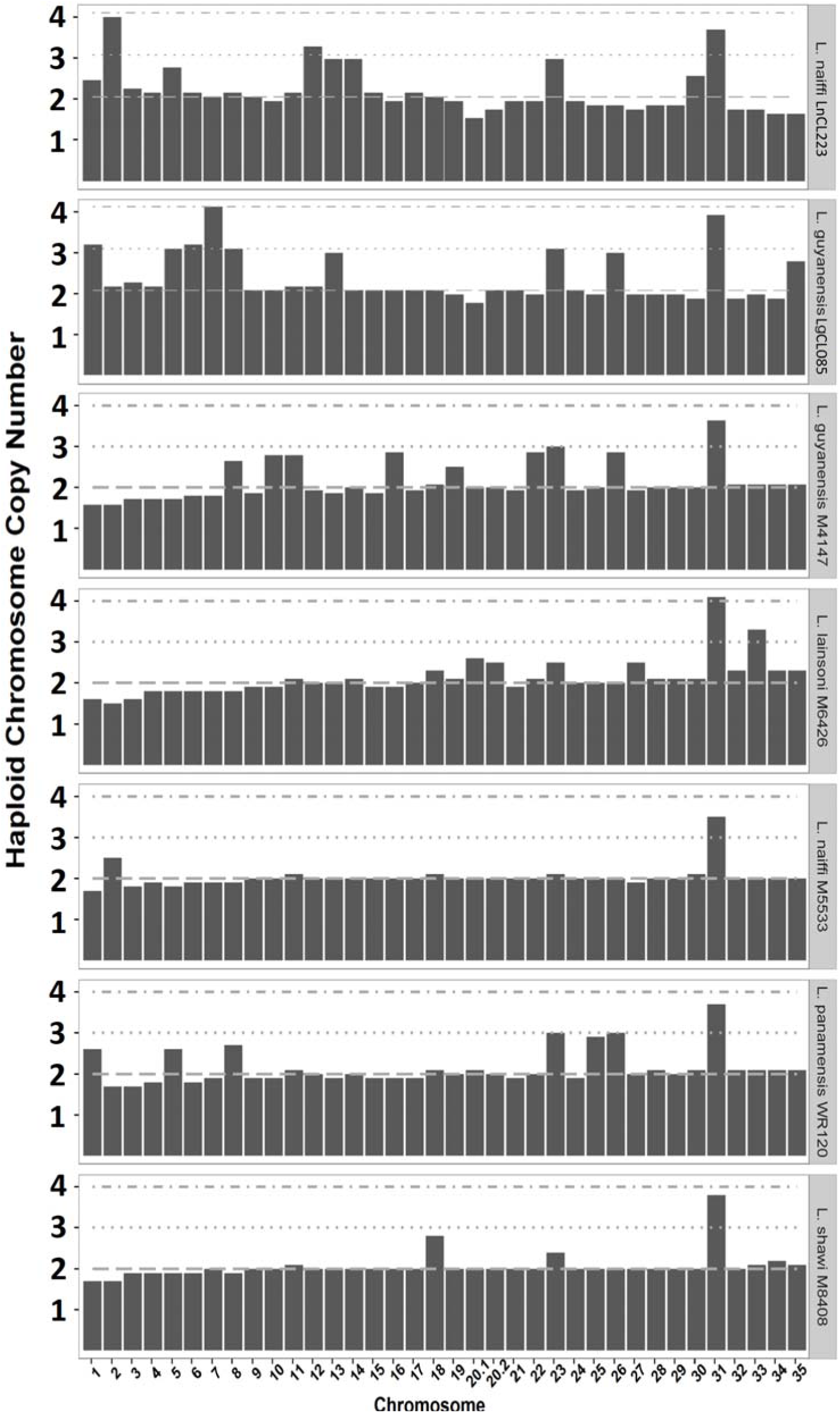
Normalised chromosome copy numbers of *L. naiffi* LnCL223 reads mapped to its assembly, *L. guyanensis* LgCL085 reads mapped to its assembly, and *L. guyanensis* M4147, *L. lainsoni* M6426, *L. naiffi* M5533, *L. panamensis* WR120 and *L. shawi* M8408 reads mapped to *L. braziliensis* M2904. Dashed lines indicate disomic, trisomic and tetrasomic states. Results for *L. panamensis* PSC-1 and *L. peruviana* PAB-4377 were previously published and are in Figure S3.

The somy patterns were supported by the results of mapping the reads of each sample to their own assembled genome or to the M2904 reference to produce the read depth allele frequency (RDAF) distributions from heterozygous SNPs. The majority of *L. braziliensis* M2904 control chromosomes had peaks with modes at ∼33% and ∼67% indicating trisomy, rather than a single peak at ∼50% consistent with disomy (Figure S5). The RDAF distributions from reads mapped to its own assembly for *L. guyanensis* LgCL085 and *L. naiffi* LnCL223 had a mode of ∼50% (Figure S6), including peaks indicating trisomy for LgCL085 chromosomes 13, 26 and 35 (Figure S7).

### 8,262 *L. naiffi* and 8,376 *L. guyanensis* genes annotated

A total of 8,262 genes were annotated on *L. naiffi* LnCL223: of these 8,104 were protein coding genes, 78 were tRNAs, 15 rRNA genes, four snoRNA genes, two snRNA genes, and 59 pseudogenes. 310 genes were on unassigned contigs (Table S3). 8,376 genes were annotated on *L. guyanensis* LgCL085: of these 8,230 were protein coding genes, 75 tRNAs, 14 rRNA genes, four snoRNA genes, two snRNA genes and 51 pseudogenes. 619 genes were on unassigned contigs

There were 8,161 genes (8,001 protein coding) transferred to the control *L. braziliensis* genome, along with 76 tRNAs, two snRNA genes, four snoRNA genes, 13 rRNA genes and 65 pseudogenes (Table 2). 7,719 of the protein coding genes (96.5%) clustered into 7,244 OGs, whereas 8,137 of the 8,375 (97.2%) protein coding genes on the *L. braziliensis* reference grouped into 7,383 OGs. This indicated that 97% of protein coding genes in OGs were recovered, and only 2.8% (235) across 201 OGs were absent in the M2904 control, mainly hypothetical or encoded ribosomal proteins (Table S5). In the same way, we found 70 protein coding genes (Table S6) in 62 OGs on the M2904 control absent in the published *L. braziliensis* annotation.

Few genes were present in *L. braziliensis* but absent in *L. guyanensis* LgCL085 and *L. naiffi* LnCL223. Coverage depth was used to predict each gene’s haploid copy number, such that genes with haploid copy numbers at least twice the assembled copy number indicated partially assembled genes in the reference assembly. Thus, we investigated all OGs with haploid copy numbers at least twice the assembled copy number to quantity completeness of the assembly. Only 145 genes in 92 OGs on *L. guyanensis* LgCL085 (Table S7), 142 genes in 90 OGs on *L. naiffi* LnCL223 (Table S8) and 102 genes in 71 OGs (Table S9) on the *L. braziliensis* control met this criterion, indicating few unassembled genes in each assembly. One hypothetical gene (LnCL223_272760) in *L. naiffi* LnCL223 with no retrievable information had a haploid copy number of 15 (OG5_173495), whereas all other genomes examined here had zero to two copies.

### A 245 Kb rearrangement akin to a minichromosome in *L. shawi* M8408

We discovered a putative minichromosome or amplification at the 3’ end of *L. shawi* M8408 chromosome 34 based on elevated coverage across a pair of inverted repeats spanning 245 Kb (Figure 3). This locus spanned at least bases 1,840,001 to 1,936,232 (the end) of *L. braziliensis* M2904 chromosome 34 (Figure S8, Table S10). It was orthologous to a known 100 Kb amplification on *L. panamensis* PSC-1 chromosome 34 that was predicted to produce a minichromosome when amplified, and contained the frequently amplified LD1 (*Leishmania* DNA 1) region [66]. In contrast to the *L. panamensis* PSC-1 minichromosome, the *L. shawi* M8408 amplification was ∼30 Kb longer and closer in length to the *L. braziliensis* M2903 245 Kb minichromosome [67].

**Figure 3:**
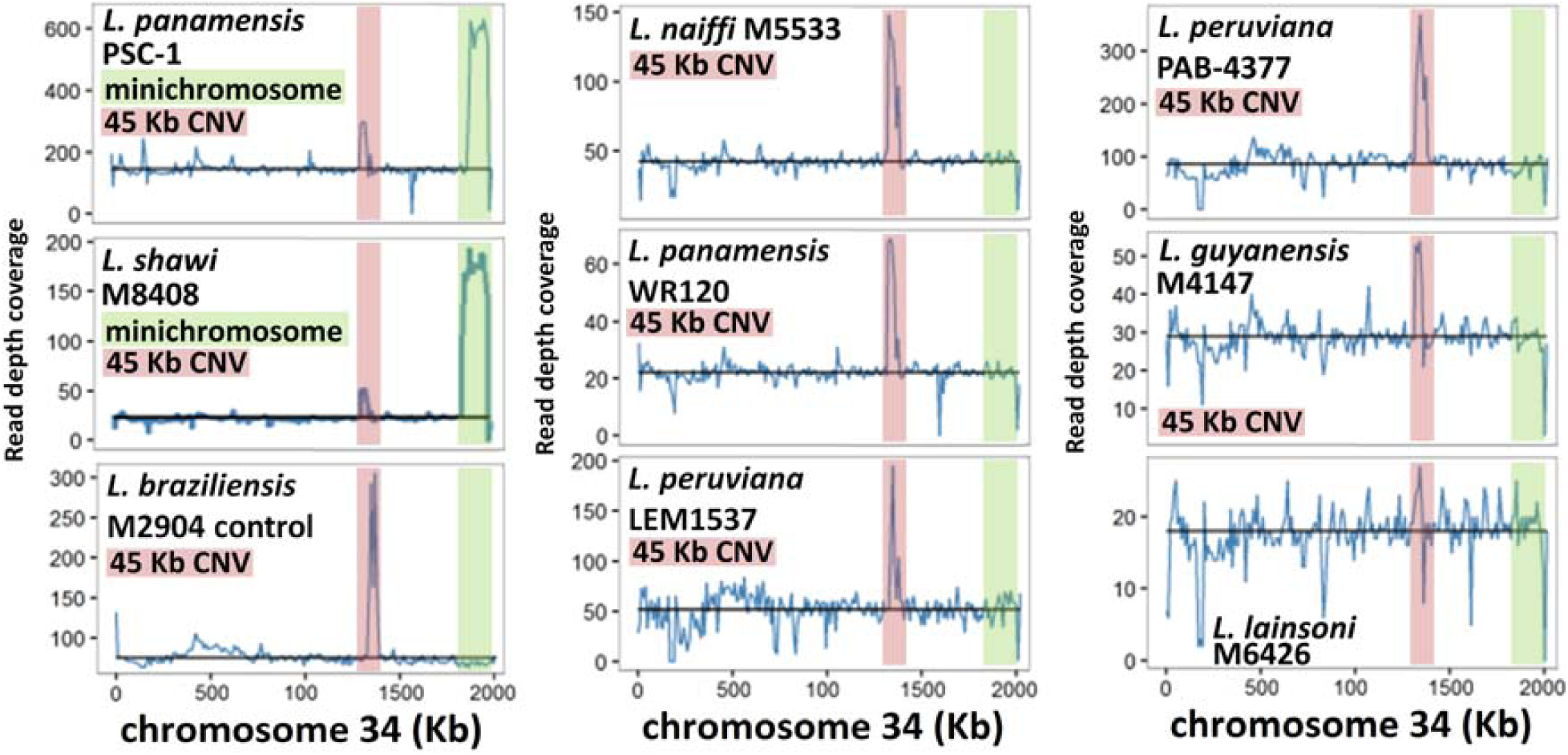
Read depth coverage (blue, y-axis) in 10 Kb blocks for reads mapped to *L. braziliensis* M2904 chromosome 34 (x-axis) for nine *Viannia* isolates. The black horizontal line is the median chromosome 34 coverage. *L. panamensis* PSC-1 (top left) and *L. shawi* M8408 (middle left) showed a 3’ jump in coverage (green) consistent with an amplification of inverted repeats that could form a linear minichromosome. In addition, this pair shared a 45 Kb amplification (pink) also found in the *L. braziliensis* M2904 control (bottom left), *L. naiffi* M5533 (top centre), *L. panamensis* WR120 (middle centre), *L. peruviana* LEM1537 (bottom centre), *L. peruviana* PAB-4377 (top right) and *L. guyanensis* M4147 (middle right). This was absent in *L. lainsoni* M6426 (bottom right).

### A 45 Kb locus was amplified in most *Viannia* genomes

A 45 Kb amplification on chromosome 34 spanning a gene encoding a structural maintenance of chromosome (SMC) family protein and ten hypothetical genes had between two and four copies in all samples except *L. lainsoni* M6426 (Figure 3, Table S10). Using the *L. guyanensis* gene annotation, putative functions were assigned to five of the ten hypothetical genes. This duplication spanned chromosomal location 1.32-1.35 Mb in the *L. braziliensis* M2904 reference and had two additional hypothetical genes in *L. naiffi* LnCL223 (LnCL223_343280 and LnCL223_343290, Figure 4).

**Figure 4:**
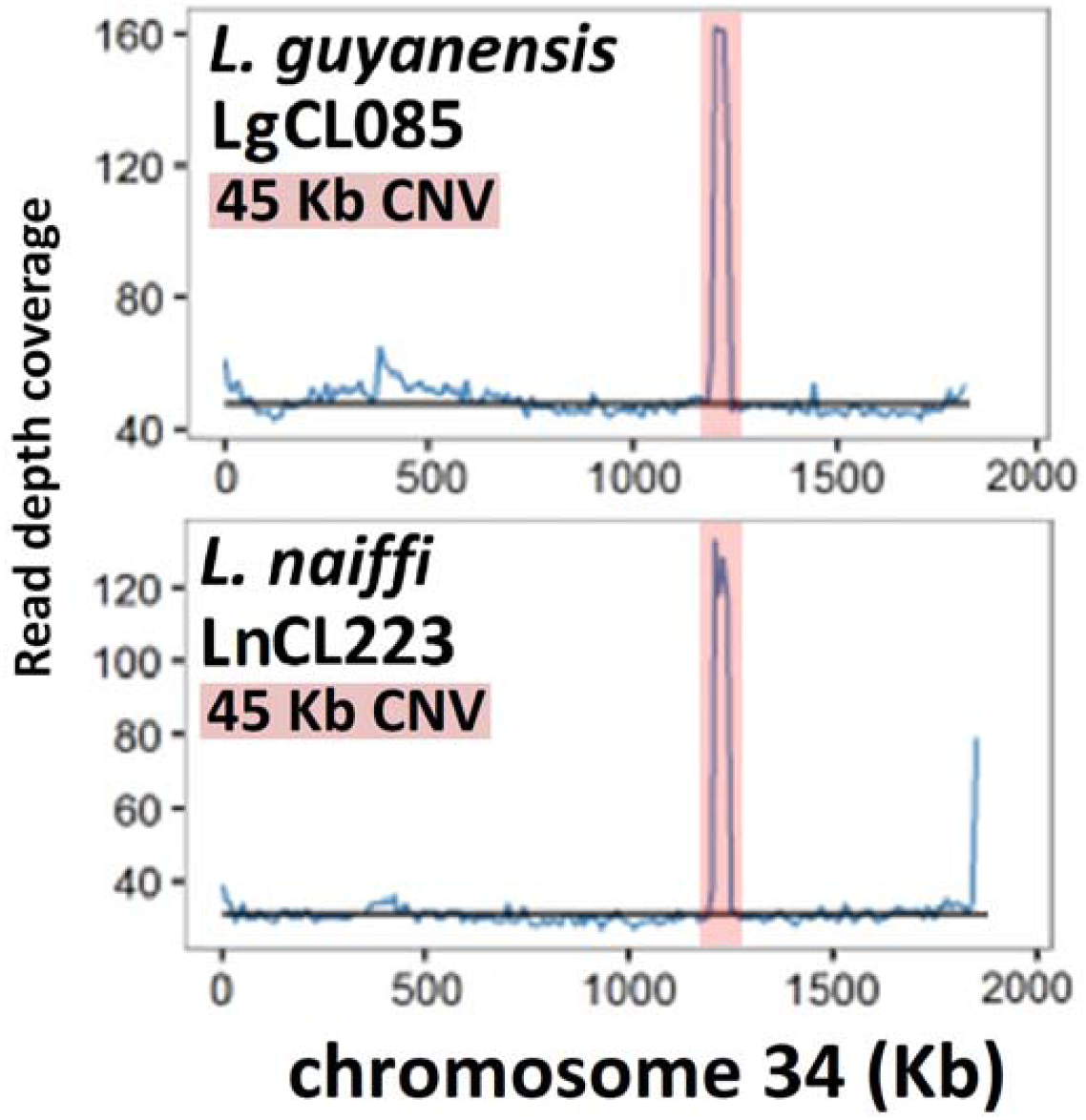
Median coverage (blue) in 10 Kb blocks for *L. guyanensis* LgCL085 reads mapped to its own assembled chromosome 34 (top) and *L. naiffi* LnCL223 reads mapped to its own assembled chromosome 34 (bottom). The black horizontal line is the median chromosome 34 read coverage. There was a 45 Kb amplification to three copies (pink) in *L. guyanensis* LgCL085 (at chromosome 34 bases 1,195,232-1,239,355, 44,123 bases in length). Similarly, there was a 45 Kb four-fold amplification (pink) in *L. naiffi* LnCL223 (at chromosome 34 bases 1,206,328-1,251,119, 44,791 bases in length). The latter encompassed two additional hypothetical genes relative to *L. guyanensis* LgCL085. Neither had evidence of a 3’ minichromosome.

### Genes exclusive to *Viannia* genomes

7,961 (96.7%) of the 8,230 genes annotated for *L. guyanensis* LgCL085 were assigned to 7,381 OGs, 7,893 (97.4%) of the 8,104 *L. naiffi* LnCL223 genes to 7,324 OGs, and 7,692 (99.3%) of the *L. panamensis* PSC-1 7,748 to 7,245 OGs. A total of 6,835 of these OGs were shared with nine species from the *Leishmania, Sauroleishmania* and *Viannia* subgenera: *L*. (*L*.) *major, L*. (*L*.) *mexicana, L*. (*L*.) *donovani* (*infantum*), *L*. (*V*.) *guyanensis, L*. (*V*.) *naiffi, L*. (*V*.) *braziliensis, L*. (*V*.) *panamensis, L*. (*S*.) *adleri, L*. (*S*.) *tarentolae* (Table S11).

We identified 22 OGs exclusive to *Viannia* (Table S12): three OGs contained the RNAi pathway genes (DCL1, DCL2, RIF4). Another OG was the telomere-associated mobile elements (TATE) DNA transposons (OG5_132061), a dynamic feature of *Viannia* genomes [51] (Supplementary Results). Four OGs encoded a diacylglycerol kinase-like protein (OG5_133291), a nucleoside transporter (OG5_134097), a beta tubulin / amastin (OG5_183241), and a /zinc transporter (OG5_214682). The remaining 14 OGs contained hypothetical genes.

A NADH-dependent fumarate reductase gene (OG5_128620) was amplified in the *Viannia* examined here: *L. guyanensis* LgCL085 had 14 copies, *L. naiffi* LnCL223 had 16, *L. panamensis* PSC-1 had 16, *L. peruviana* PAB4377 had 23, *L. peruviana* LEM1537 had 14, and *braziliensis* M2904 had 12. This contrasted with the *Leishmania* and *Sauroleishmania* subgenera for which three to four copies had been reported for *L. infantum, L. mexicana, L. major, L. adleri* and *L. tarentolae* [68,69]. This gene has been implicated in enabling parasites to resist oxidative stress and potentially aiding persistence, drug resistance and metastasis [70,71].

### Few species-specific genes in *L. guyanensis* LgCL085 and *L. naiffi* LnCL223

Four genes from four OGs unique to *L. naiffi* LnCL223 were identified compared to other *Leishmania* (Table S13). Of these four, hypothetical genes LnCL223_312570 and LnCL223_292920 had orthologs in *T. brucei* and *T. vivax*, respectively. The LnCL223_341350 protein product had 44-45% sequence identity with a *Leptomonas* transferase family protein, and LnCL223_352070 was a methylenetetrahydrofolate reductase (OG5_128744), but had no orthologs in the other eight *Leishmania* or five *Trypanosoma* species investigated here. *L. guyanensis* LgCL085 had 31 unique genes in 30 OGs, 25 of which were on unplaced contigs. Four of the six chromosomal genes were also in *Trypanosoma* genomes, encoding two hypothetical proteins (a tuzin and a poly ADP-ribose glycohydrolase). 28 of the 31 had orthologs in eukaryotes, of which three had orthologs in the free-living freshwater ciliate protozoan *Tetrahymena thermophile* (Table S14) [72].

### *L. guyanensis* LgCL085 and *L. naiffi* LnCL223 had over 300 gene arrays

Gene arrays are genes in the same OG with more than two haploid gene copies: they can be *cis* or *trans*. There were 327 gene arrays on *L. naiffi* LnCL223 (Table S15), 334 on *L. guyanensis* LgCL085 (Table S16) and 255 on the control *L. braziliensis* M2904 (Table S17) – half the arrays on each genome had two copies of each gene. 22 of the *L. guyanensis*, 18 of the *L. naiffi* LnCL223 and 15 of the control *L. braziliensis* gene arrays contained 10+ haploid gene copies (Table 3). The *L. panamensis* PSC-1 genome had ∼400 tandem arrays, of which 71% had more than two copies. The *L. braziliensis* M2904 genome had 615 arrays corresponding to 763 OGs in OrthoMCL v5. Thus, the control genome underestimated the number of gene arrays due to either gene absence or incomplete assembly, indicating that the number of arrays on *L. naiffi* LnCL223 and *L. guyanensis* LgCL085 was underestimated.

**Table 3:** Arrays with ten or more gene copies predicted by read depth for each species. OG stands for orthologous group. Genes in OG shows the number of genes associated with that GO functional category. OG haploid copy number indicates the numbers of haploid gene copies found in each genome: B stands for the *L. braziliensis* M2904 control, G for *L. guyanensis* LgCL085, and N for *L. naiffi* LnCL223. *For OG5_126623 and OG5_126588, the elevated copy number were due to amplificated heat shock protein (*hsp*) genes rather than the glucose regulated protein (*grp*) loci, a potential limitation of OG analyses.

| OG |  | Genes in OG |  |  | OG haploid copy number |  |  |
| --- | --- | --- | --- | --- | --- | --- | --- |
| ID | Description | B | G | N | B | G | N |
| ***L. guyanensis* LgCL085, *L. naiiffi* LnCL223 and *L. braziliensis* M2904 control** |  |  |  |  |  |  |  |
| OG5_132061 | TATE DNA Transposon | 2 | 14 | 3 | 21 | 50 | 11 |
| OG5_126605 | alpha tubulin | 1 | 2 | 1 | 17 | 36 | 13 |
| OG5_130729 | amastin-like surface protein | 8 | 24 | 20 | 24 | 26 | 33 |
| OG5_126631 | elongation factor 1-alpha | 1 | 1 | 1 | 12 | 18 | 20 |
| OG5_126703 | polyubiquitin | 2 | 2 | 1 | 21 | 16 | 43 |
| OG5_126558 | dynein heavy chain, cytosolic | 14 | 13 | 14 | 13 | 13 | 14 |
| OG5_129265 | pteridin transporter; folate/biopterin transporter | 8 | 10 | 9 | 14 | 13 | 13 |
| OG5_143904 | amastin-like surface protein | 5 | 4 | 6 | 48 | 12 | 14 |
| OG5_126623* | lipophosphoglycan biosynthetic protein / glucose regulated protein 94; heat shock protein 90 / 83-1 | 2 | 2 | 2 | 11 | 10 | 12 |
| ***L. guyanensis* LgCL085 and *L. naiiffi* LnCL223** |  |  |  |  |  |  |  |
| OG5_126749 | GP63 leishmanolysin | 4 | 8 | 9 | 5 | 33 | 56 |
| OG5_126617 | receptor-type adenylate cyclase | 3 | 5 | 5 | 5 | 14 | 13 |
| OG5_126611 | beta tubulin | 1 | 1 | 2 | 0 | 14 | 27 |
| OG5_128620 | NADH-dependent fumarate reductase | 4 | 3 | 4 | 2 | 14 | 16 |
| OG5_126585 | kinesin K39; hypothetical protein | 7 | 10 | 10 | 7 | 12 | 11 |
| OG5_126573 | histone H4 | 1 | 7 | 4 | 3 | 11 | 10 |
| ***L. naiiffi* LnCL223** |  |  |  |  |  |  |  |
| OG5_173495 | hypothetical protein | 0 | 1 | 1 | 0 | 2 | 15 |
| OG5_126568 | ABC1 transporter | 9 | 10 | 12 | 10 | 9 | 14 |
| OG5_127342 | peptidase m20/m25/m40 family protein | 2 | 2 | 2 | 6 | 3 | 10 |
| ***L. guyanensis* LgCL085** |  |  |  |  |  |  |  |
| OG5_173452 | tuzin | 2 | 3 | 1 | 1 | 19 | 1 |
| OG5_145872 | ATG8/AUT7/APG8/PAZ2 | 1 | 2 | 1 | 8 | 19 | 1 |
| OG5_143922 | ATP dependent DEAD-box helicase | 1 | 2 | 0 | 6 | 17 | 0 |
| OG5_148241 | conserved hypothetical protein | 1 | 1 | 1 | 0 | 14 | 1 |
| OG5_137181 | ATG8/AUT7/APG8/PAZ2 | 1 | 1 | 0 | 0 | 13 | 0 |
| ***L. braziliensis* M2904 control** |  |  |  |  |  |  |  |
| OG5_126588* | heat-shock protein hsp70; glucose-regulated protein 78 | 4 | 3 | 3 | 14 | 7 | 8 |
| OG5_138994 | tuzin | 5 | 3 | 2 | 13 | 4 | 2 |
| OG5_129839 | phosphoglycan beta 1,3 galactosyltransferase | 2 | 4 | 1 | 12 | 5 | 2 |
| OG5_127067 | thimet oligopeptidase; metallo-peptidase, Clan MA(E), Family M3 | 3 | 3 | 3 | 12 | 9 | 7 |
| ***L. guyanensis* LgCL085 and *L. braziliensis* control** |  |  |  |  |  |  |  |
| OG5_169610 | surface antigen-like protein | 1 | 1 | 0 | 16 | 11 | 0 |
| OG5_127518 | SLACS like gene retrotransposon element | 2 | 3 | 1 | 18 | 10 | 1 |

The most expanded array on *L. guyanensis* LgCL085 contained TATE DNA transposons (OG5_132061) with 50 haploid gene copies (Table 3) compared with 11 on *L. naiffi* LnCL223, 21 on the *L. braziliensis* control and 16 on *L. panamensis* PSC-1. The *L. braziliensis* M2904 assembly had 40 TATE DNA transposons, but only two were annotated on the control here, illustrating that more accurate estimates of copy number may be possible.

*L. naiffi* LnCL223 had the highest haploid gene copy number of the M8 family metalloprotease leishmanolysin (GP63) array (OG5_126749) with 56 haploid gene copies, compared to 33 in *L. guyanensis* LgCL085, 28 in *L. panamensis* PSC-1 and 31 in *L. braziliensis* M2904. This was the sole protease-related OG amplified in all three species (Table S23). This family was not expanded in *L. peruviana* LEM1537 or PAB4377. This was consistent with previous work on *L. guyanensis* leishmanolysin [73] indicating it is a highly expressed virulence factor in promastigotes [74] affecting the survival during the initial stages of infection [74-77]. *Sauroleishmania* genomes also had high array copy numbers: 37 for *L. adleri* [69] and 84 for *L. tarentolae* (Table S12). *Leishmania* subgenus genomes had lower copy numbers, with 13 for *L. mexicana*, 15 for *L. infantum* and five for *L. major* (OG4_10176 for *L. braziliensis* M2904, *L. mexicana, L. infantum* and *L. major*).

A tuzin gene array (OG5_173452) had higher haploid copy numbers on *L. guyanensis* LgCL085 (19) and *L. panamensis* PSC-1 (22) compared with the two copies in *L. naiffi, L. mexicana, L. infantum, L. major, L. braziliensis, L. adleri* and *L. tarentolae*. Tuzins are conserved transmembrane proteins in *Trypanosoma* and *Leishmania* associated with surface glycoprotein expression [78]. They are often contiguous with δ-amastin genes, whose products are abundant cell surface transmembrane glycoproteins potentially involved in the infection or survival within macrophages. They are absent in *Crithidia* and *Leptomonas* species, who lack a vertebrate host stage [78]. Tuzins may play a role in pathogenesis [79], which may be related to leishmaniasis caused by *L. guyanensis*.

## Discussion

### *L. (Viannia) guyanensis* and *L. (V*.*) naiffi* draft reference genomes

We assembled high-quality reference genomes for two isolates, *L. (Viannia) guyanensis* LgCL085 and *L. (V*.*) naiffi* LnCL223, from short read sequence libraries to illuminate genomic diversity in the *Viannia* subgenus and extend previous work [52]. This process combined the *de novo* assembly with a reference-guided approach using the published genome of *L. braziliensis* M2904 to assemble the *L. guyanensis* LgCL085 and *L. naiffi* LnCL223 into 35 chromosomes each (Table 2). An essential feature of this process was to identify and remove contamination in the *L. guyanensis* and *L. braziliensis* M2904 libraries and to trim low-quality bases in *L. naiffi* LnCL223 to ensure that the reads used were informative and free of exogenous impurities. A second screen for contamination in unassigned contigs also removed several *L. guyanensis* LgCL085 contigs, which improved subsequent annotation and gene copy number estimates.

### Genomes assembled from short reads capture aneuploidy and nearly all genes

Our strategy was tested by applying the same protocol to the *L. braziliensis* M2904 short read library, which acted as a positive control and quantified the precision of the final output. This facilitated the detection of structural variation or annotation problems, chiefly underestimated copy numbers at certain genes and the incorrect assembly of some loci that were fixed manually. The resulting genomes were largely complete: for comparison, the control *L. braziliensis* M2904 genome had only four homozygous SNPs, 97.2% of the protein coding genes of the reference (231 were missing) and 70 additional genes missed in the reference sequence. These findings highlight scope to resolve *Leishmania* chromosomal architecture more accurately, particularly at repetitive regions and gene arrays, using longer sequencing reads and hybrid assembly approaches.

We showed that the majority of *Viannia* were diploid and had 35 chromosomes. Aneuploidy was evident for *L. guyanensis* LgCL085, *L. guyanensis* M4147, *L. naiffi* LnCL223, *L. naiffi* M5533, *L. lainsoni* M6426, *L. panamensis* WR120 and *L. shawi* M8408 as anticipated [80]. This was verified using read depth allele frequency distributions of reads mapped to *L. braziliensis* M2904 and to their own assemblies.

The *L. guyanensis* LgCL085 genome had more protein coding genes (8,230) than *L. naiffi* LnCL223 (8,104). These numbers were similar to those for *L. panamensis* PSC-1 (7,748) [51] and *L. braziliensis* M2904 (8,357) [48]. The vast majority of protein coding gene models were computationally transferred [81] from the *L. braziliensis* M2904 reference with perfect matching, and were verified and improved manually. Both the *L. guyanensis* and *L. naiffi* reference genomes contained unassigned bin contigs, and chromosomal regions homologous to multiple chromosomal loci or containing partially collapsed gene arrays. 90 (*L. naiffi*) and 92 (*L. guyanensis*) collapsed gene arrays were identified where haploid gene copy numbers were at least twice the assembled copy number when the reads were mapped to the assembled genomes.

### A better resolution of the *Viannia* species complexes

This study illustrated that high-throughput sequencing approaches, alignment methods and annotation tools can improve the accuracy of *Leishmania* gene copy number estimates, gene organistion, and genome structure resolution. This yielded insights into features differentiating the isolates examined here, including a 45 Kb duplication on chromosome 34 of most *Viannia*, variable gene repertoires across *Viannia* species, and a potential minichromosome derived from the 3’ end of *L. shawi* M8408 chromosome 34. Further work is required to investigate *L. utingensis* and *L. lindenbergi* and other potential distinct lineages [82].

Both single-gene and large-scale copy number variations (CNVs) were tolerated by all *Leishmania* genomes. *Leishmania* genomes have extensive conservation of gene content with few species-specific genes [45,48]: here, only 31 *L. guyanensis* LgCL085 and four *L. naiffi* LnCL223 species-specific genes were found. These four genes unique to *L. naiffi* LnCL223, its leishmanolysin hyper-amplification, the 31 genes only in *L. guyanensis* LgCL085 and its tuzin arrays all represent potential targets for improving species-specific typing and better disease surveillance. This is important because infections by the *Viannia* are spread by many hosts and all sources of infections need to be addressed. Immunological screening of anti-*Leishmania* antibodies could be enhanced by genetic testing to identify infections from non-endemic or rarer sources like *L. naiffi*, which has longer parasite survival rates in macrophages *in vitro* [83].

MLSA of 100 *Viannia* isolates across four genes and genome-wide diversity inferred from mapped reads indicated that *L. guyanensis* LgCL085 was closest to *L. panamensis* PSC-1 within the *L. guyanensis* species complex, but was assigned the *L. guyanensis* classification because *L. guyanensis, L. panamensis* and *L. shawi* were a monophyletic species complex as shown by MLSA [56], MLMT [64], *hsp70* [65], internal transcribed spacer (ITS) [84,85], MLEE [86] and RAPD data [87]. Further typing of a more extensive *L. guyanensis, L. panamensis* and *L. shawi* isolate set might clarify if these are distinct species or a single genetic group.

More precise genetic screening of *Viannia* isolates is necessary to trace hybridisation between species. Infection of humans, dogs and *Lu. ovallesi* with *L. guyanensis/L. braziliensis* hybrids was reported in Venezuela [40-41]. A *L. shawi/L. guyanensis* hybrid causing CL was detected in Amazonian Brazil [42], and *L. naiffi* has produced viable progeny with *L. lainsoni* [43] and *L. braziliensis* (Elisa Cupolillo, unpublished data). There is extensive evidence of interbreeding among *L. braziliensis* complex isolates, including more virulent *L. braziliensis/L. peruviana* hybrids with higher survival rates within hosts *in vitro* [44].

## Conclusion

This study highlighted the utility of genome sequencing for the identification, characterisation and comparison of *Leishmania* species. We demonstrated that short reads were sufficient for assembly of most *Leishmania* genomes so that SNP, chromosome copy number, structural and somy changes can be investigated comprehensively. The *L. (Viannia) guyanensis* and *L. (V*.*) naiffi* genomes represent a further advance in refining the taxonomical complexity of the *Viannia* by illustrating their genomic characteristics and the extent to which these are shared across *Viannia* species, which will assist examining the extent to which they can hybridise. This improved understanding of *Leishmania* genomes should be used to explore the complex epidemiology of CL and MCL pathologies in the Americas and the roles of non-human reservoirs and sand flies in these processes. Future work could tackle transmission, drug resistance and pathogenesis in the *Viannia* by applying long-read high-throughput sequencing to examine broader sets of isolates, their genetic diversity, contributions to microbiome variation, and control of transcriptional dosage at gene amplifications.

## Methods

### *L. guyanensis* and *L. naiffi* whole genome sequencing

Extracted DNA for *L. guyanensis* LgCL085 and *L. naiffi* LnCL223 was received from Charité University Medicine (Berlin) at the Wellcome Trust Sanger Institute on 6th Feb 2012. Paired-end 100 bp read Illumina HiSeq 2000 libraries were prepared for both during which *L. guyanensis* required 12 cycles of PCR. The DNA was sequenced (run 7841_5#12) on the 15^th^ (*L. guyanensis*, run 7841_5#12) and 23^rd^ (*L. naiffi*, run 7909_7#9) March 2012. The library preparation, sequencing and read quality verification was conducted as outlined previously [69]. The resulting *L. guyanensis* library contained 15,272,969 reads with a median insert size of 327.0 (NCBI accession ERX180458) and the *L. naiffi* one had 8,131,246 reads with a median insert size of 335.4 (ERX180449).

### *Viannia* comparative genome, annotation and proteome files

The *L. braziliensis* reference genome (MHOM/BR/1975/M2904) was a positive control whose short reads were examined using the same methods. It was originally sequenced using an Illumina Genome Analyzer II [48] yielding 26,007,384 76 bp paired-end reads with a median insert size of 244.1 bp (ERX005631). Protein sequences were retrieved from the EMBL files using Artemis [88]. Two *L. panamensis* genomes, two *L. peruviana* genome assemblies and five 100 bp paired-end Illumina HiSeq 2000 read libraries of other *Viannia* isolates [53] were used for comparison (Table 1). We included the genomes of *L. panamensis* MHOM/PA/1994/PSC-1, *L. peruviana* PAB-4377 and LEM1537 (MHOM/PE/1984/LC39), and the 100 bp Illumina HiSeq 2000 paired-end reads for each *L. peruviana* PAB-4377 (16,117,316 reads) and *L. peruviana* LEM1537 (9,378,317 reads).

### Library quality control, contaminant removal and screening

Figure S9 presents an overview of the bioinformatic steps used in this paper. Quality control of the *L. guyanensis* LgCL085, *L. naiffi* LnCL223, *L. braziliensis* M2904, the five *Viannia* libraries from [53], two *L. peruviana* libraries and *L. panamensis* PSC-1 read library was carried out using FastQC (www.bioinformatics.babraham.ac.uk/projects/fastqc/). No corrections were required for the other libraries. An abnormal distribution of GC content per read observed as an extra GC content peak outside the normal peak for the *L. braziliensis* M2904 and *L. guyanensis* reads indicated sequence contamination that was removed (Figure S10). Two Illumina PCR primers in the *L. braziliensis* M2904 reads were removed (Table S1). Further evaluation using GC content filtering and the non-redundant nucleotide database with BLASTn [89] to remove contaminant sequences (Figure S10) with subsequent correction of read pairing arrangements reduced the initial 52,014,768 reads to 34,592,618 properly paired reads for assembly.

The M2904 reads used to assemble a control genome were used for read mapping, error correction and SNP calling and so the contamination did not affect the published reference. However, it did reduce the number of reads mapped as shown in [48] where only 84% of the *L. braziliensis* M2904 short reads mapped to the *L. braziliensis* assembly, compared with 92% of reads for *L. infantum* reads mapped to its own assembly, 93% of *L. major* reads mapped to its own assembly, and 97% of *L. mexicana* reads mapped to its own assembly.

The 8,131,246 100 bp paired-end *L. naiffi* LnCL223 reads and 15,272,969 100 bp paired-end *L. guyanensis* LgCL085 reads were filtered (Table S1) in the same manner using BLASTn and the smoothness of the GC content distribution to remove putative contaminants. Low quality bases were trimmed at the 3’ end of *L. naiffi* LnCL223 reads to remove bases with a phred base quality < 30 using Trimmomatic [90] (Table S1, Figure S11). This resulted in 13,033,846 paired-end *L. guyanensis* LgCL085 sequences and 6,989,814 paired-end *L. naiffi* LnCL223 sequences – 85% and 86% of the initial reads, respectively (Table S1).

### Genome evaluation, assembly and optimisation

Processed reads were assembled into contigs using Velvet v1.2.09 and assemblies for all odd numbered k-mer lengths from 21 to 75 were evaluated. The expected k-mer coverage was determined for each assembly using the mode of a k-mer coverage histogram from the velvet-estimate-exp_cov.pl script in Velvet to maximise resolution of repetitive and unique regions [57]. This suggested optimal k-mers of 61 for *L. guyanensis* LgCL085 and 43 for both *L. naiffi* LnCL223 and *L. braziliensis*, which produced assemblies with the highest N50 lengths. Each assembly was assembled with this expected coverage, and contigs were removed if their average k-mer coverage was less than half the expected coverage levels. An expected coverage of 16 and a coverage cutoff of 8 was applied to *L. naiffi* reads, an expected coverage of 19 and coverage cutoff of 8.5 to *L. guyanensis* LgCL085, and an expected coverage of 28 and coverage cutoff of 14 to *L. braziliensis*.

The assembly with the highest N50 for each was scaffolded using SSPACE [58]. In the initial assemblies, 76% of gaps in scaffolds (3,592/4,754) were closed in for *L. guyanensis* LgCL085, 63% (4,096/6,530) for *L. naiffi* LnCL223, and 67% (4,834/8,786) for *L. braziliensis* using Gapfiller [58]. Erroneous bases were corrected by mapping reads to the references with iCORN [91] (Figure S12). Misassemblies detected and broken using REAPR [60] were aligned to the *L. braziliensis* M2904 reference (excluding the bin chromosome 00). Scaffolds were evaluated and broken at putative misassemblies detected from the fragment coverage distribution (FCD) error and regions with low coverage when the reads were mapped to both broken and unbroken options. Additionally, the *L. braziliensis* broken and unbroken scaffolds were used to verify that removing misassemblies prior to (but not after) the contiguation of scaffolds resulted in more accurate assembled chromosomes. Mis-assembled regions without a gap were replaced with N bases. REAPR corrected 444 errors in *L. naiffi* LnCL223, of which 59 were caused by low fragment coverage, 206 in *L. guyanensis* LgCL085 (eight due to low fragment coverage), and 232 in the *L. braziliensis* control (57 caused by low fragment coverage). Each assembly step improved the corrected N50 and percentage of error free bases (EFB%) assessed using REAPR (Table S18), with the sole exception of *L. braziliensis* control at the error-correction stage, likely due to its higher heterozygosity. The EFB% was the fraction of the total bases whose reads had no mismatches, matches the expected insert length, had a small FCD error and at least five read pairs oriented in the expected direction.

Gaps > 100 bp were reduced to 100 bp. 200 bp at the edge of each unplaced scaffold was aligned with the 200 bp flanking all pseudo-chromosome gaps using BLASTn to verify that no further gaps could be closed using unplaced scaffolds. Unplaced bin scaffolds < 1 Kb were discarded, and the resulting assemblies were visualised and compared to *L. braziliensis* using the Artemis Comparison Tool (ACT) [92]. *L. guyanensis* LgCL085 bin sequences with BLASTn E-values < 1e-05 and percentage identities > 40% to non-*Leishmania* species in non-redundant nucleotide database were removed as possible contaminants. The final scaffolds were contiguated using the *L. braziliensis* reference with ABACAS [59], unincorporated segments were labelled as unassigned “bin” contigs, and kDNA contigs were annotated as well (Supplementary Methods).

### Phylogenomic MLSA characterisation

A MLSA (multi-locus sequence analysis) approach was adopted to verify the *Leishmania* species identity using for four housekeeping genes: glucose-6-phosphate dehydrogenase (G6PD), 6-phosphogluconate dehydrogenase (6PGD), mannose phosphate isomerase (MPI) and isocitrate dehydrogenase (ICD). Orthologs from other genomes and assemblies were obtained using BLASTn alignment with thresholds of E-value < 0.05 and percentage identity > 70%. *L. peruviana* LEM-1537 genome had gaps at the MPI and 6PGD genes and was excluded. The four housekeeping genes spanning 2,902 sites were concatenated in the order G6PD, 6PGD, MPI and ICD, and aligned using Clustal Omega v1.1 to create a Neighbour-Net network of uncorrected p-distances using SplitsTree v4.13.1.

### Genome annotation and manual curation

Annotation of the *L. guyanensis* LgCL085, *L. naiffi* LnCL223 and *L. braziliensis* control genomes was completed using Companion [80] using *L. braziliensis* M2904 as the reference as outlined previously [69], including manual checking and correction of gene models. A control run with the *L. braziliensis* M2904 reference genome using itself as a reference was performed. In *L. naiffi* LnCL223, 13 genes and one pseudogene were removed because they overlapped existing superior gene models that had improved sequence identity with *L. braziliensis* M2904 orthologs. 46 of the protein coding genes were also manually added. 34 of the protein coding genes on *L. guyanensis* LgCL085 were manually added and one protein coding gene was removed. 269 gene models on *L. naiffi* LnCL223 and 198 on *L. guyanensis* with multiple joins mainly caused by the presence of short gaps were corrected by extending the gene model across the gap where the gap length was known (< 100 bp). If the gap length was unknown (> 100 bp), the gene was extended to the nearest start or stop codon.

### Measuring ploidy, chromosome copy numbers and CNVs

By mapping the reads with SMALT v5.7 (www.sanger.ac.uk/resources/software/smalt/) to *L. braziliensis* M2904, the coverage at each site was determined to quantify the chromosome copy numbers and RDAF distributions at heterozygous SNPs as per previous work [69]. The RDAF distribution was based on the coverage level of each allele at heterozygous SNPs and this feature differed across chromosomes for each isolate (Supplementary Results). The median coverage per chromosome was obtained, and the median of the 35 values combined with the RDAF distribution mode approximating 50% indicated that all isolates examined here were mostly diploid (except the triploid *L. braziliensis* M2904). These were visualised with R packages ggplot2 and gridExtra.

After PCR duplicate removal, the mapped reads were used to detect CNVs across genes or within non-overlapping 10 Kb blocks for all chromosomes and bin contigs using the median depth values normalised by the median of the chromosome (or bin contig). Loci with a copy number > 2 were analysed for *L. naiffi* LnCL223, *L. guyanensis* LgCL085 and the *L. braziliensis* control using their reads mapped to their own assembly. This was also repeated for reads mapped to the *L. braziliensis* M2904 reference for *L. guyanensis* M4147, *L. naiffi* M5533, *L. shawi* M8408, *L. lainsoni* M6426, *L. panamensis* WR120, *L. panamensis* PSC-1, *L. peruviana* LEM1537 and *L. peruviana* PAB-4377. *L. panamensis* PSC-1 reads were mapped to its own reference genome to verify that we could find previously identified amplified loci, and we mapped *L. panamensis* WR120 to it so that CNVs shared by both *L. panamensis* could be obtained. The BAM files of *L. naiffi* LnCL223, *L. guyanensis* LgCL085 and *L. braziliensis* M2904 reads mapped to its own assembly were visualised in Artemis to confirm and refine the boundaries of amplified loci.

### Identification of orthologous groups and gene arrays

Protein coding genes from *L. guyanensis* LgCL085, *L. naiffi* LnCL223 and the *L. braziliensis* M2904 control genome were produced from the EMBL files for each genome and these were submitted to the ORTHOMCLdb v5 webserver [93] to identify orthologous groups (OGs).

11,825 OGs with associated gene IDs in at least one of four *Leishmania* species (*L. major* strain Friedlin, *L. infantum, L. braziliensis* and *L. mexicana)* or five *Trypanosoma* species (*T. vivax, T. brucei, T. brucei gambiense, T. cruzi* strain CL Brener and *T. congolense*) were retrieved from the OrthoMCL database and compared with OGs for each genome. The copy number of each OG was estimated by summing the haploid copy number of each gene in the OG. Gene arrays in each genome were identified by finding all OGs with haploid copy number > 2. Large arrays (> 10 gene copies) were examined and arrays with unassembled gene copies were identified by finding those with haploid gene copy number at least twice the assembled gene number.

### SNP screening and detection

The filtered reads with Smalt as described mapped above were used for calling SNPs using Samtools Pileup v0.1.11 and Mpileup v0.1.18 and quality-filtered with Vcftools v0.1.12b and Bcftools v0.1.17-dev as previously [69] such that SNPs called by both Pileup and Mpileup post-screening were considered valid. These SNPs all had: base quality >25; mapping quality >30; SNP quality >30; a non-reference RDAF >0.1; forward-reverse read coverage ratios >0.1 and <0.9; five or more reads; 2+ forward reads, and 2+ reverse reads. Low quality and repetitive regions of the assemblies were identified and variants in these regions were masked as outlined elsewhere [69]. SNPs were classed as homozygous for an alternative allele to the reference if their RDAF > 0.85 and heterozygous if it was > 0.1 and < 0.85.

The high level of nucleotide accuracy of the assembled genomes was indicated by the low rate of homozygous SNPs when the reads mapped to its own assembly (50 for *L. naiffi* LnCL223, 12 for *L. guyanensis* LgCL085, 68 for the *L. braziliensis* reference, and four for the *L. braziliensis* control). Likewise, the numbers and alleles of heterozygous SNPs for the *L. braziliensis* control (25,474) matched that for the reference (25,975), suggesting that the 705 (*L. naiffi* LnCL223) and 14,739 (*L. guyanensis* LgCL085) heterozygous SNPs were accurate. The difference in homozygous and heterozygous SNP rates for *L. braziliensis* here versus the original 2011 study [48] were likely due to differing methodology. The genetic divergence of *L. naiffi* LnCL223 and *L. guyanensis* LgCL085 compared to *L. braziliensis* was quantified using the density of heterozygous and homozygous SNPs per 10 Kb non-overlapping window on each chromosome, visualised using Bedtools.

## Data Accessbility

The BioProject ID is PRJEB20208 for *L. guyanensis* LgCL085 and PRJEB20209 for *L. naiffi* LnCL223. The DNA reads are available at the NCBI Short Read Archive (SRA) and and European Nucleotide Archive at ERX180458 for *L. guyanensis* LgCL085 and ERX180449 for *L. naiffi* LnCL223 (these are associated with BioProject PRJEB2600). The consensus genome sequence FASTA files are on Figshare at 10.6084/m9.figshare.5693290 for *L. guyanensis* LgCL085 and 10.6084/m9.figshare.5693272 for *L. naiffi* LnCL223. The chromosome and bin contig annotation EMBL files are at 10.6084/m9.figshare.5693284 for *L. guyanensis* LgCL085 and 10.6084/m9.figshare.5693278 for *L. naiffi* LnCL223. The Supplementary Tables are on Figshare at 10.6084/m9.figshare.5697064. For ease of reader access, the above genome sequence and annotation files, Supplementary Tables and Supplementary Data are also available on the Dryad Digital Repository at: https://doi.org/10.5061/dryad.4bm23.

## Acknowledgements

The authors thank: Matthew Berriman and members of the WTSI DNA pipelines team for generating the two sequence libraries; Elisa Cupolillo (Instituto Oswaldo Cruz, Brazil) for discussions and comments on the manuscript; Katrin Kuhls (Technical University of Applied Sciences Wildau), Cathal Seoighe (NUI Galway), Hideo Imamura and Jean-Claude Dujardin (both Institute of Tropical Medicine Antwerp) for help; Anne Stone and Kelly Harkins (both Arizona State University) for releasing valuable sequence read data; and the DJEI/DES/ SFI/HEA Irish Centre for High-End Computing (ICHEC) for computational facilities.

## Contributions

S.C. completed the genome assembly, comparative genomics, phylogenetic analysis, mutation investigation, helped design the study and wrote the main manuscript text. S.C., A.S.T. and E.F. completed the genome annotation. M.S. completed genome sequencing. G.S. helped design the study and wrote the main manuscript text. J.A.C. helped design the study and wrote the main manuscript text. T.D. co-ordinated and designed the study and wrote the main manuscript text. All authors gave approval for publication.

