## Supplementary Materials for "*Leishmania naiffi* and *Leishmania guyanensis* reference genomes highlight genome structure and gene evolution in the *Viannia* subgenus"

**Supplementary Data**

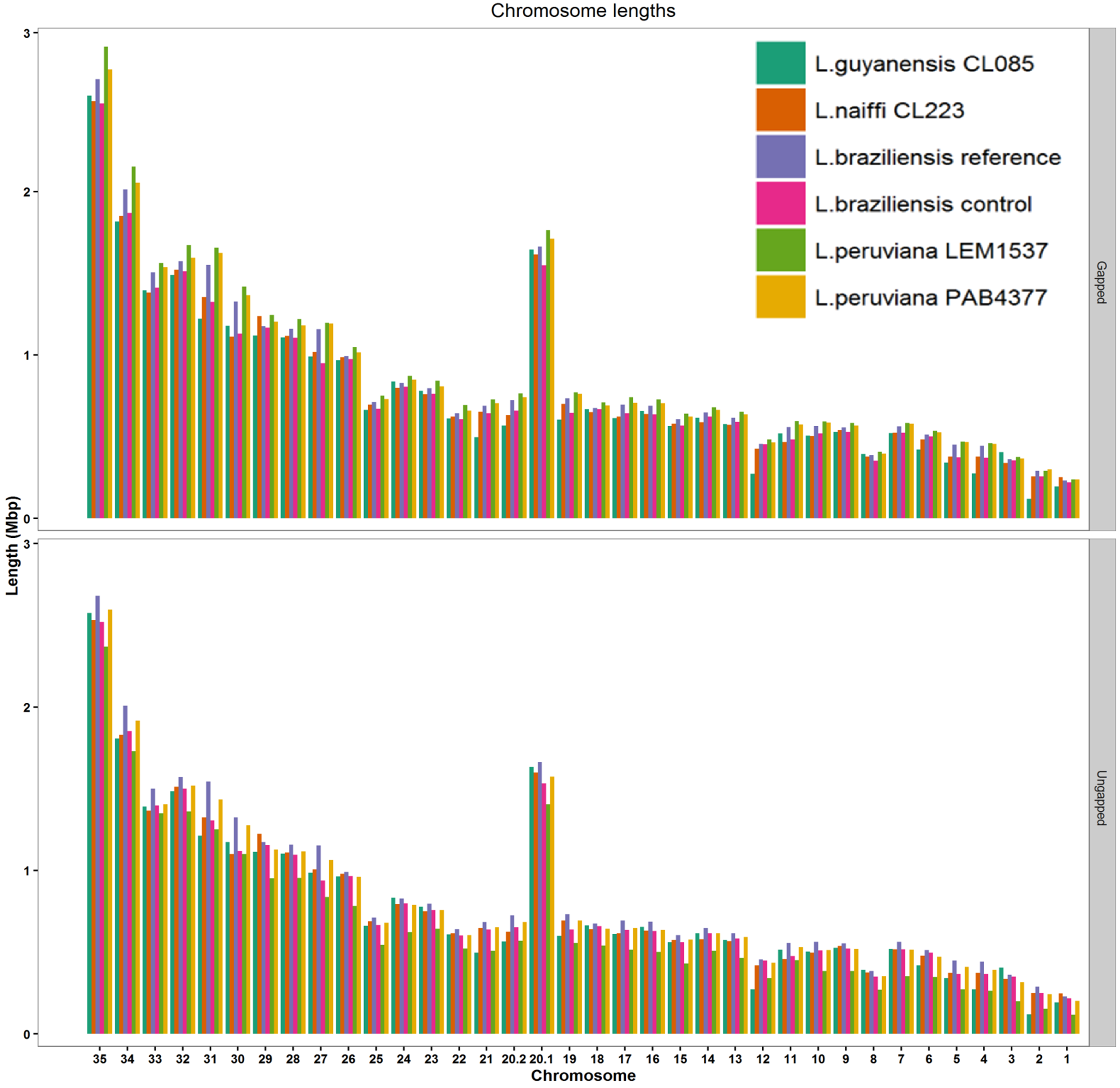

**Figure S1**: Chromosome lengths of *L. guyanensis* LgCL085, *L. naiffi* LnCL223 and the *L. braziliensis* M2904 control compared with the *L. braziliensis* M2904 reference genome, *L. peruviana* PAB-4377, *L. peruviana* LEM1537 and *L. panamensis* PSC-1. Lengths were examined both including gaps in the chromosome length (top) and excluding gaps from chromosome lengths (bottom).

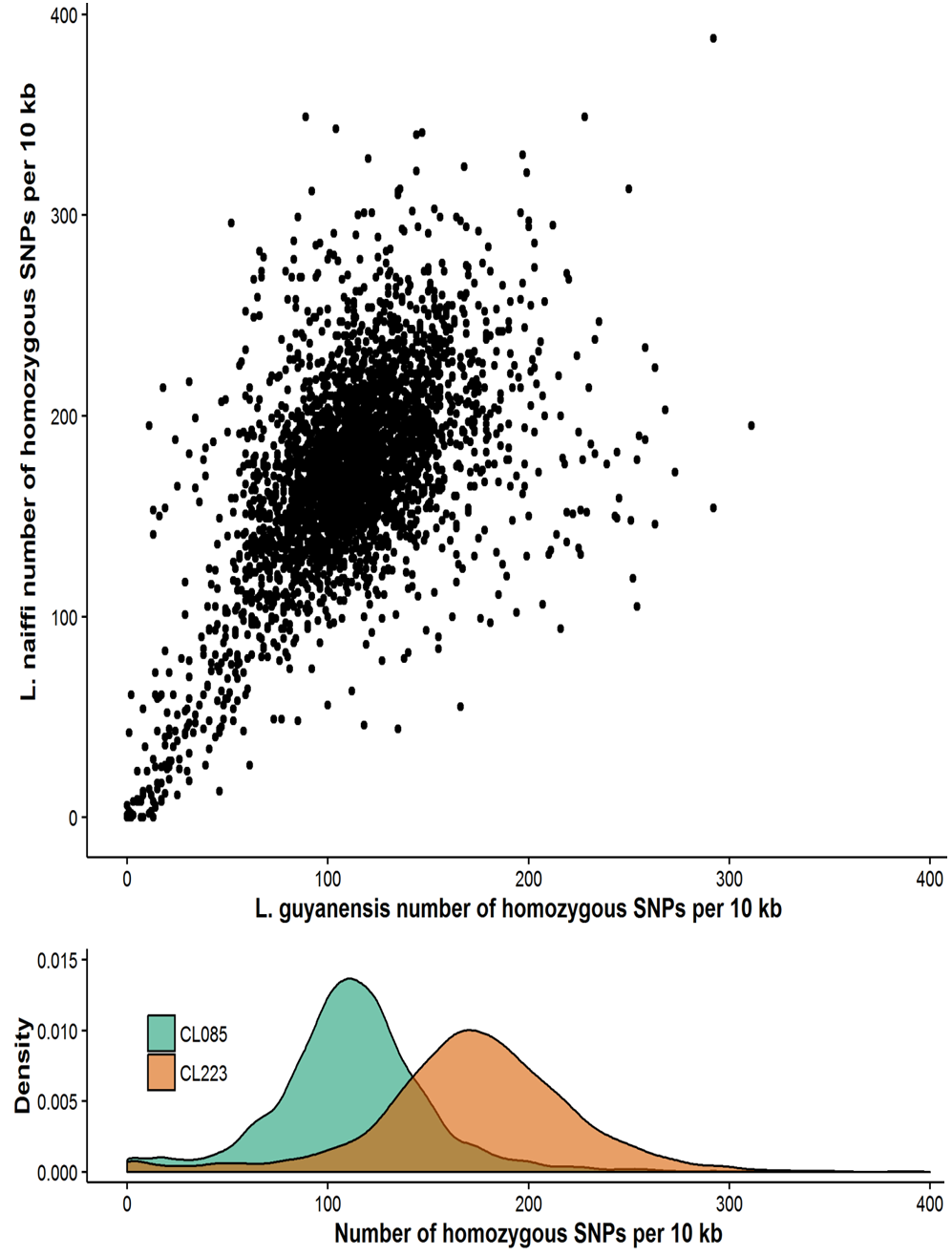

**Figure S2:** The genome-wide rate of homozygous SNPs per non-overlapping 10 Kb block shown (Top panel) for *L. guyanensis* LgCL085 reads (x-axis, green in lower panel) and *L. naiffi* LnCL223 reads (y-axis, orange in lower panel) showed no evidence of of differential evolutionary rates nor recent gene flow, though (Bottom panel) *L. guyanensis* was more closely related to *L. braziliensis* M2904 (green) than *L. naiffi* LnCL223 (orange).

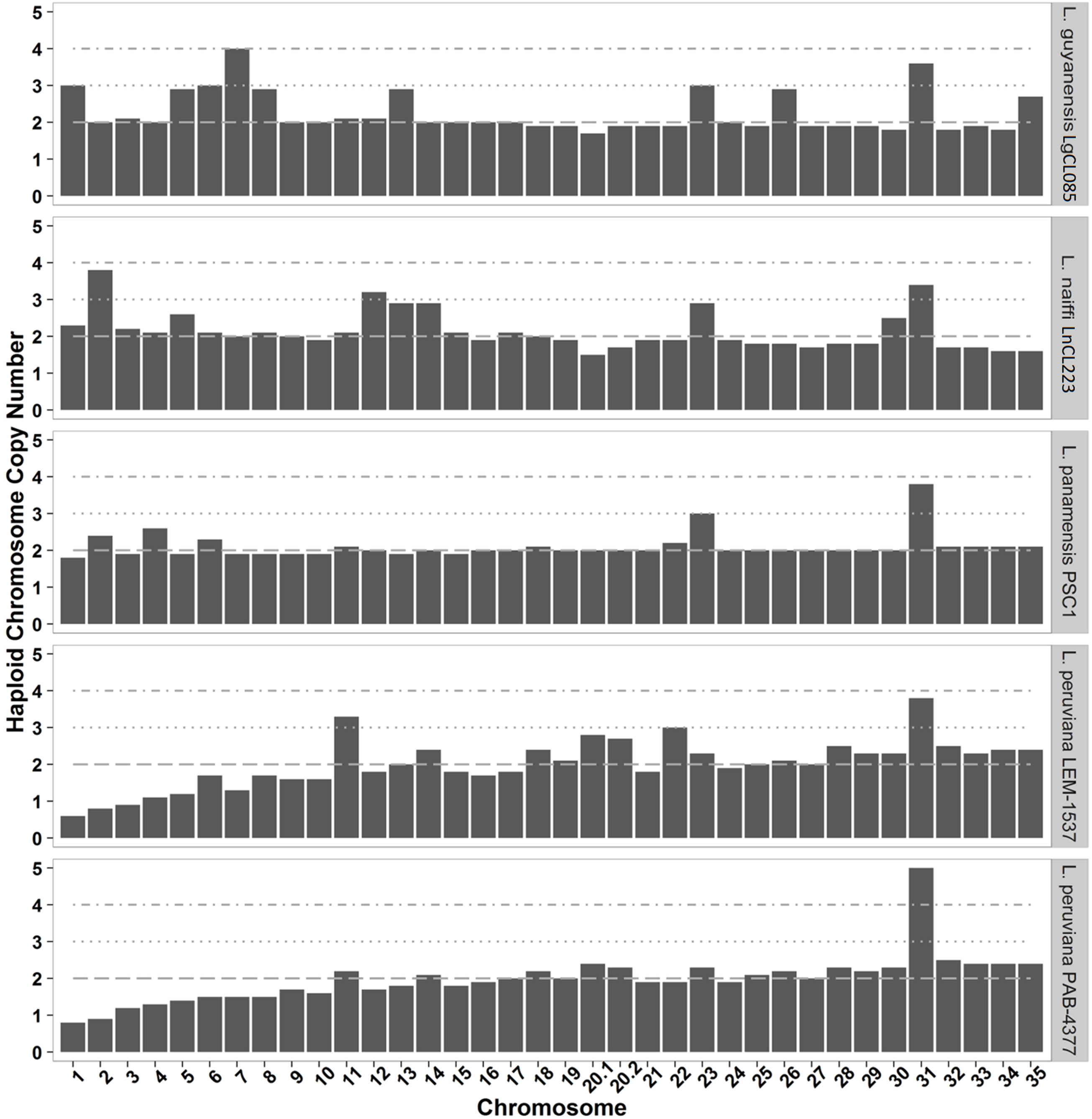

**Figure S3**: Chromosome copy numbers predicted using coverage of reads mapped to *L. braziliensis* M2904 for: *L. guyanensis* LgCL085, *L. naiffi* LnCL223, *L. panamensis* PSC-1, *L. peruviana* LEM-1537 and *L. peruviana* PAB-4377.

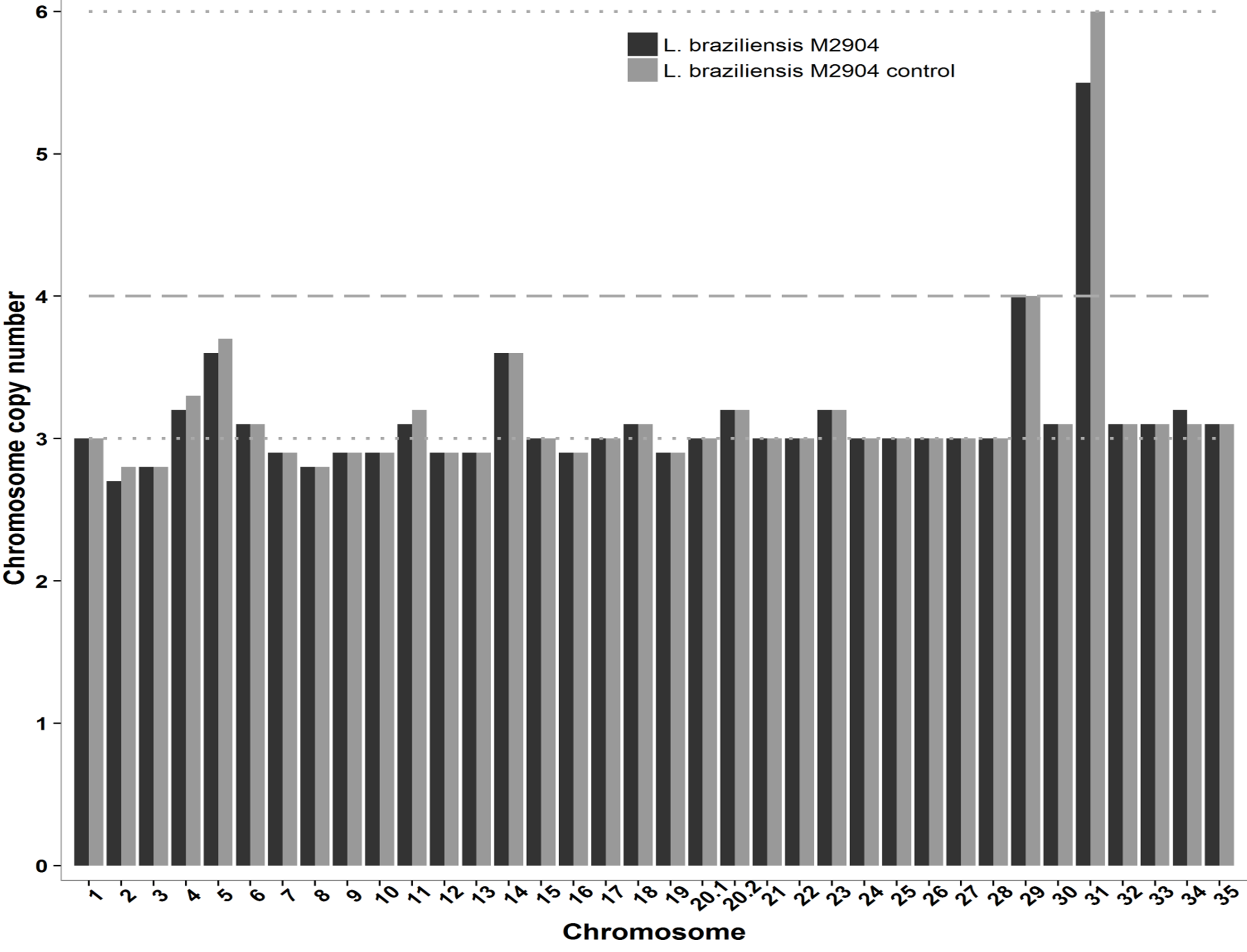

Figure S4: Chromosome copy numbers of the control *L. braziliensis* M2904 assembly: this replicated the results of Rogers et al, 2011 for *L. braziliensis* M2904.

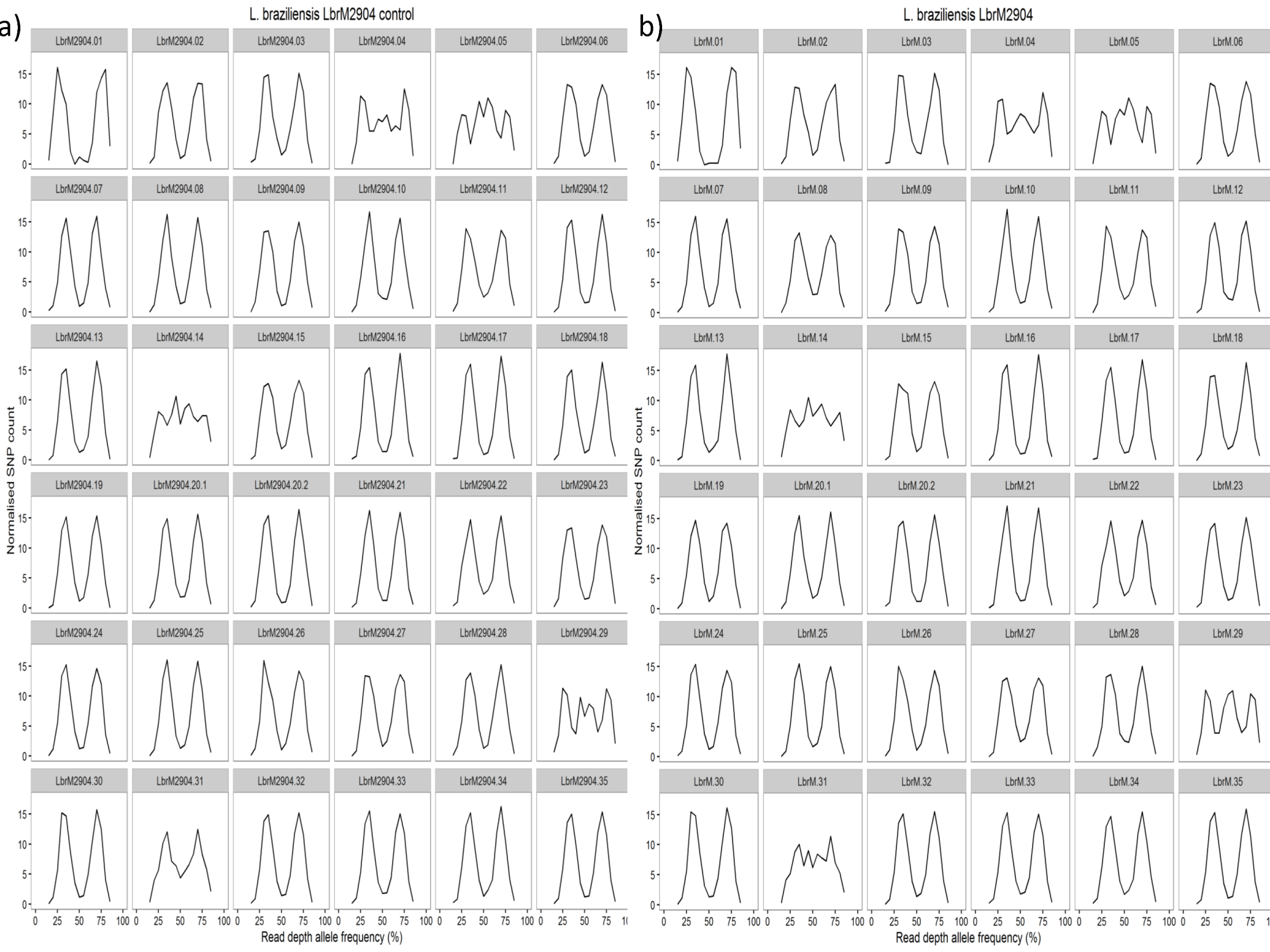

Figure S5: Read depth allele frequency distributions (RDAF) of heterozygous SNPs called from reads mapped to its own assembly for: a) each chromosome of *L. braziliensis* M2904 control genome, compared with b) each chromosome of the reference *L. braziliensis* M2904 genome. This showed that both produce the same distributions for every chromosome.

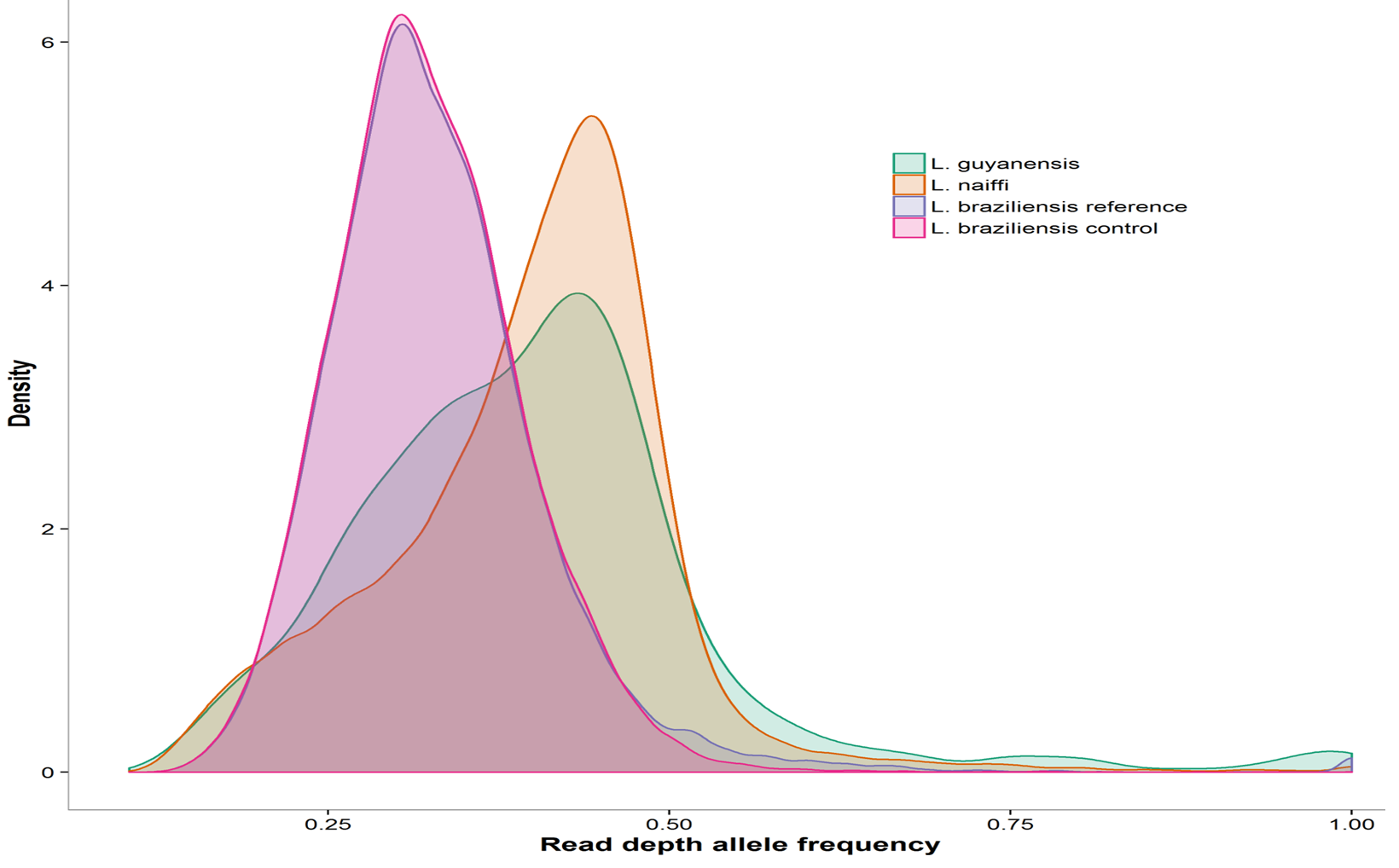

Figure S6: Read depth allele frequencies (RDAF) of SNPs for *L. naiffi* LnCL223 reads mapped to its own assembled genome, *L. guyanensis* LgCL085 reads mapped to its own assembled genome, and the *L. braziliensis* reads mapped to its reference genome as well as the control assembled genomes examined here. This showed that *L. naiffi* LnCL223 and *L. guyanensis* LgCL085 were mainly disomic but *L. braziliensis* was largely trisomic.

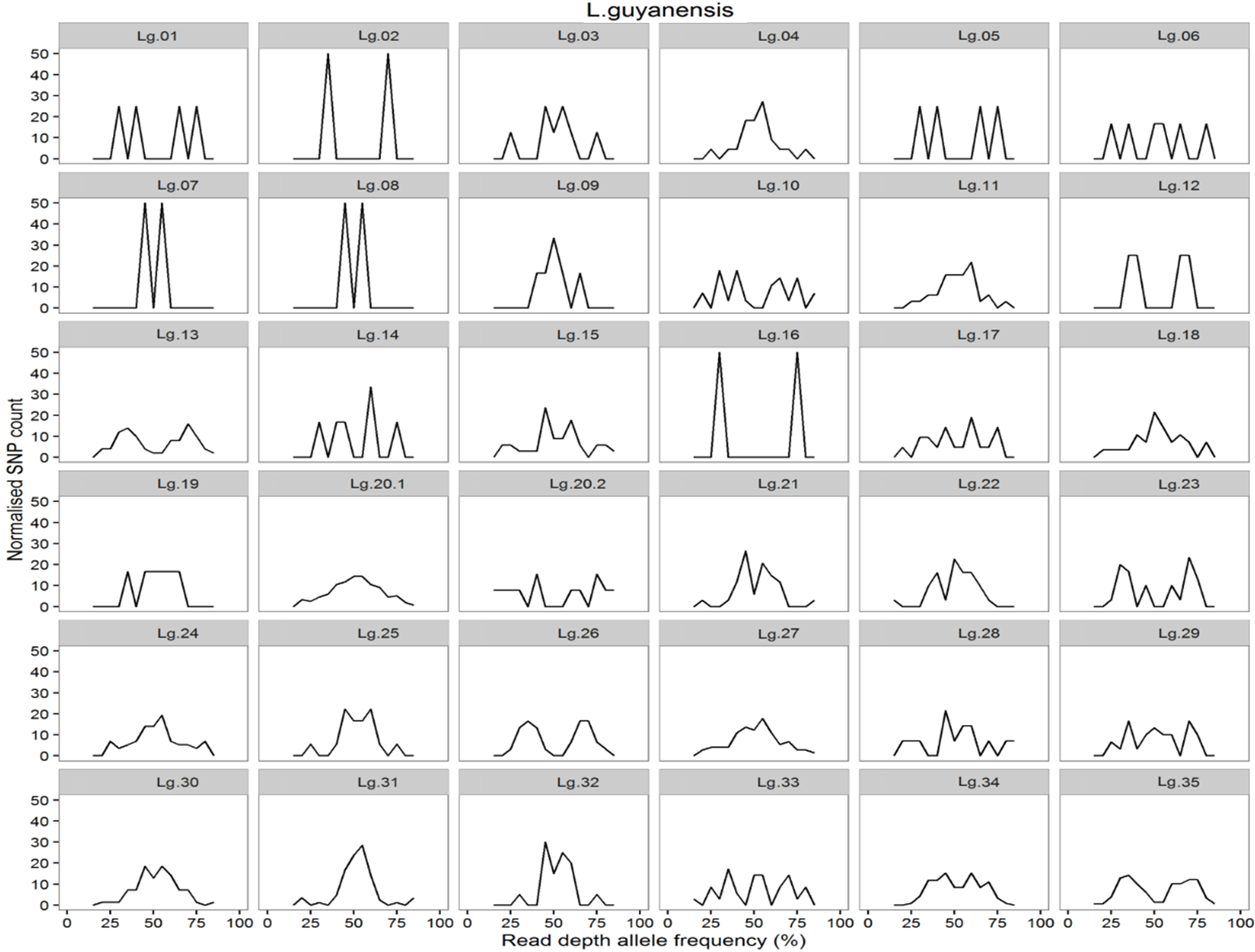

Figure S7: Read depth allele frequency distributions (RDAF) of each *L. guyanensis* LgCL085 chromosome determined using heterozygous SNPs from reads mapped to its own assembly. Most chromosomes had uninformative RDAF plots here due to low number of heterozygous SNPs.

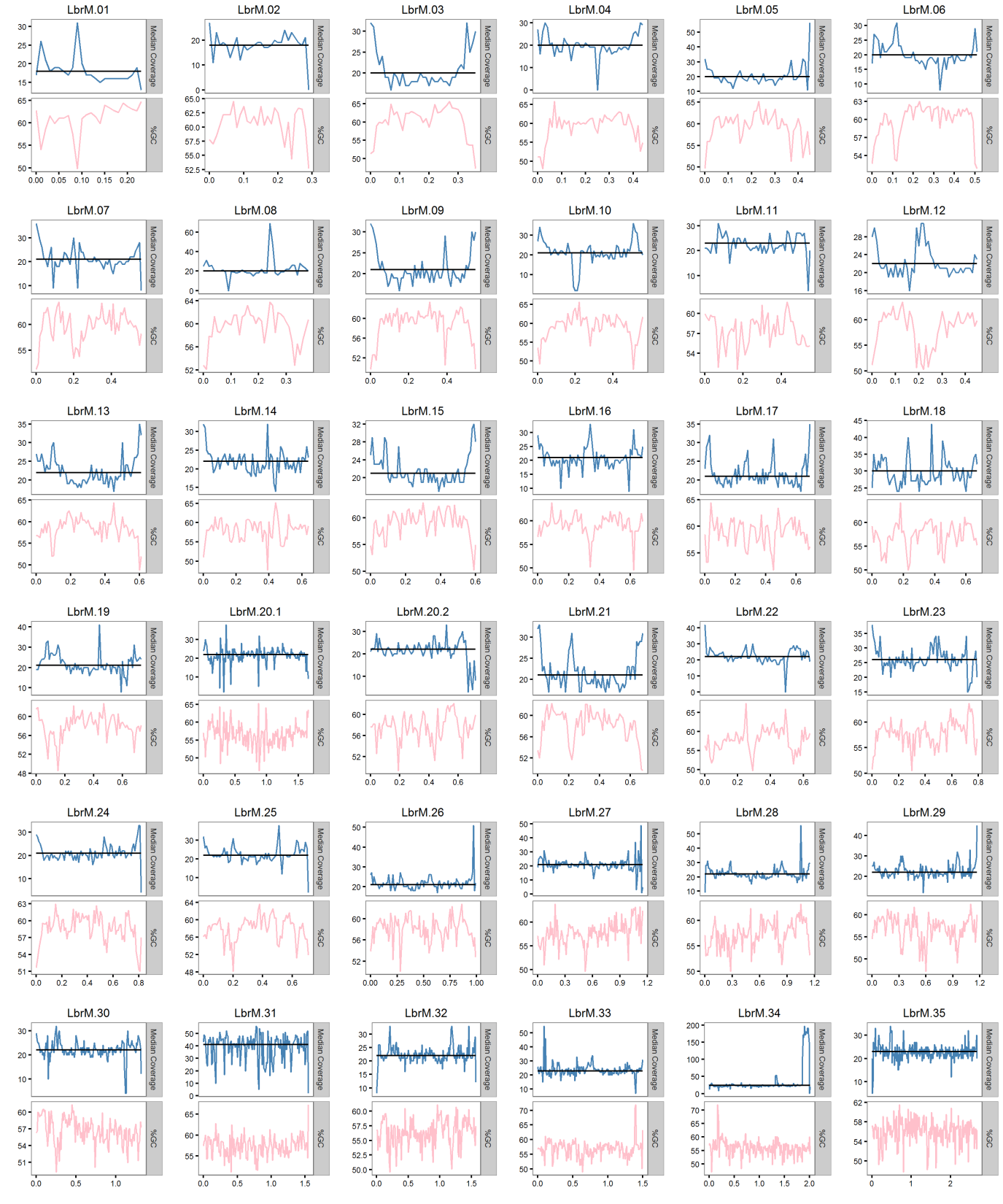

Figure S8: Median coverage (blue) in 10 kb blocks across each chromosome for *L. shawi* M8408 reads mapped to *L. braziliensis* M2904. The x-axis units are Mb. The black horizontal line in each plot denotes the median chromosomal coverage and the pink line indicates %GC content measured in 10 kb blocks. Note the increase in coverage at the 3’ end of chromosome 34 (second last plot on the bottom row) indicating amplification of inverted repeats at that locus that could form a linear minichromosome.

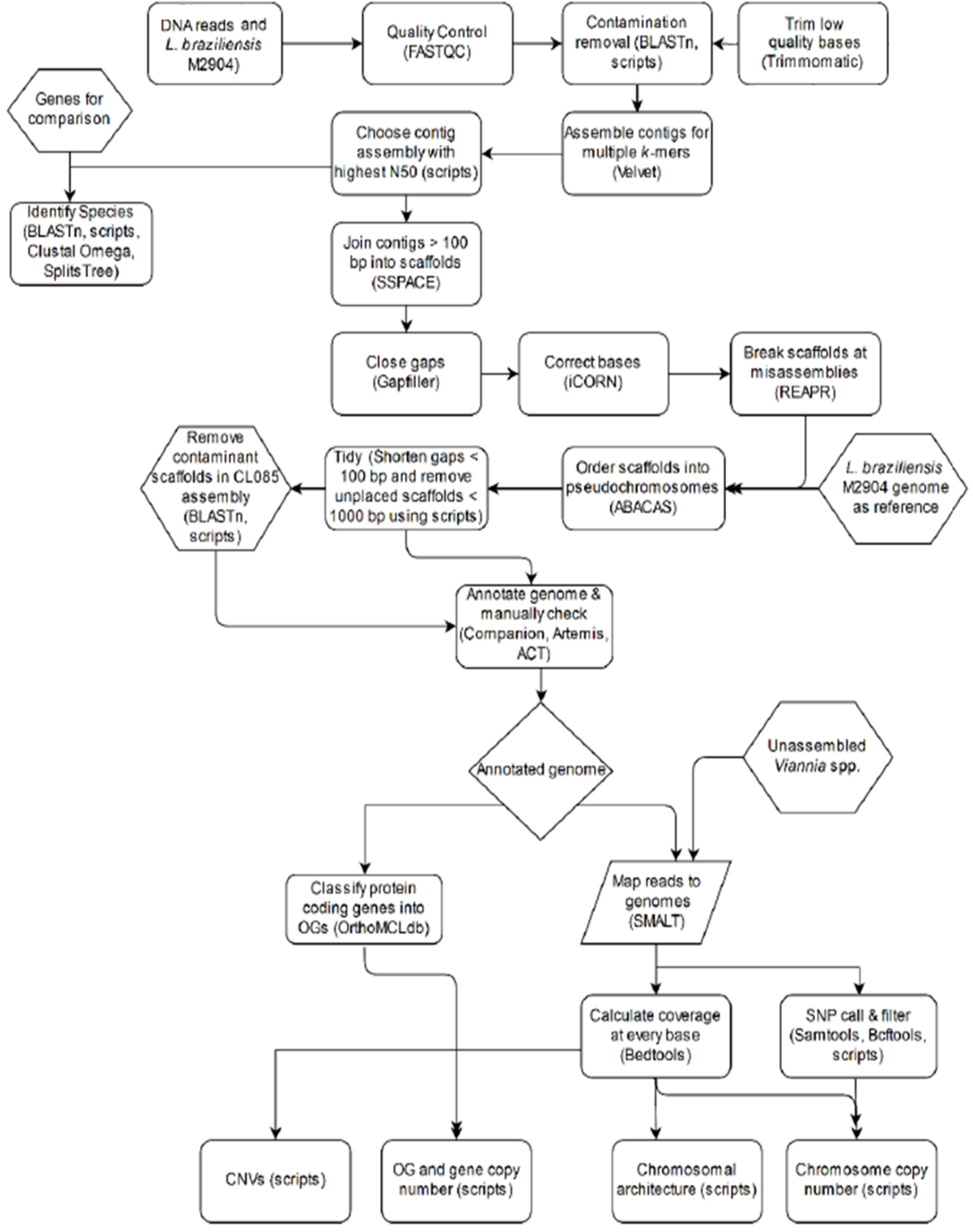

**Figure S9:** A graphical summary of *L. naiffi* LnCL223 and *L. guyanensis* LgCL085 genome assembly, improvement, annotation and analysis.

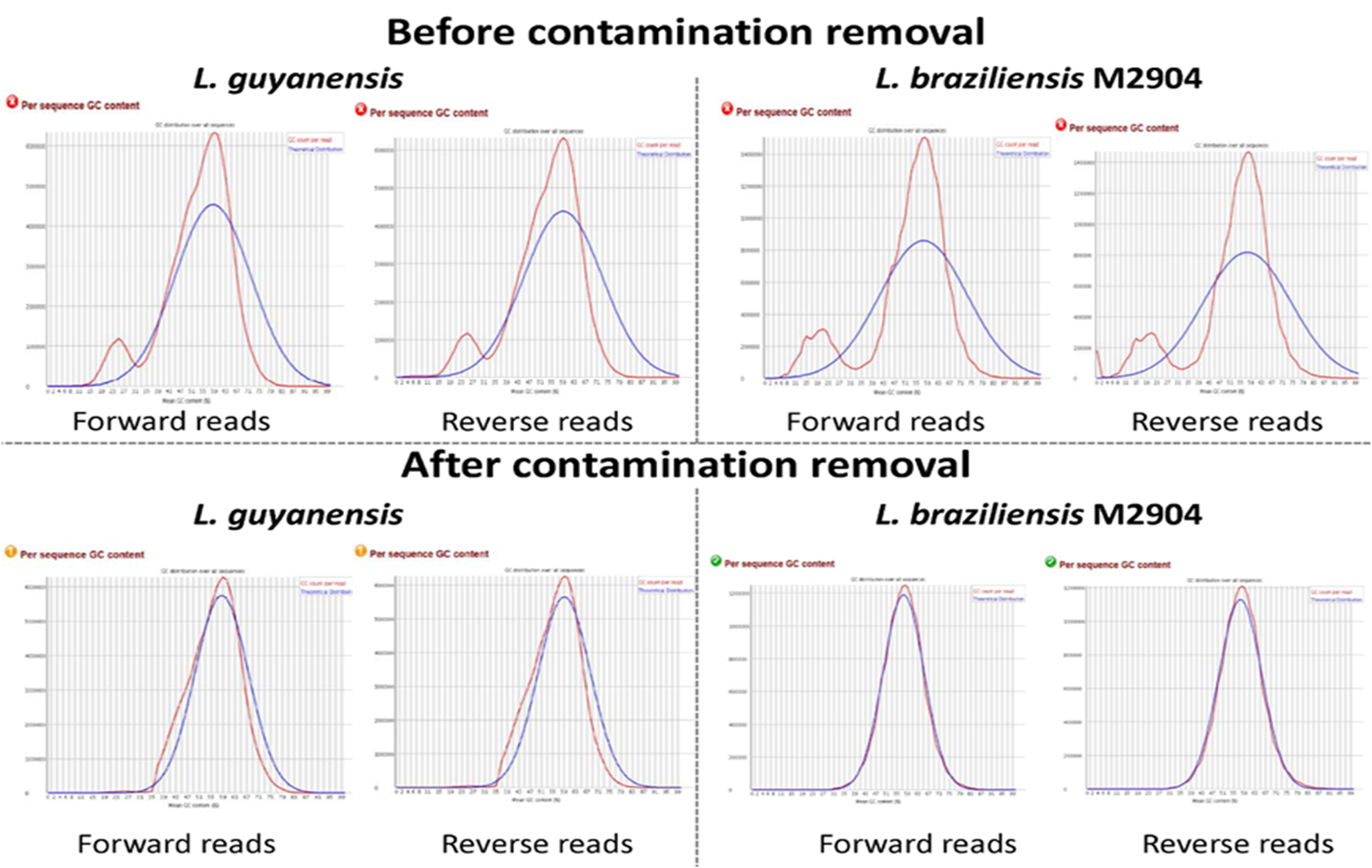

Figure S10: GC content plots produced by FASTQC of *L. guyanensis* LgCL085 and *L. braziliensis* M2904 Illumina short reads before and after contamination removal.

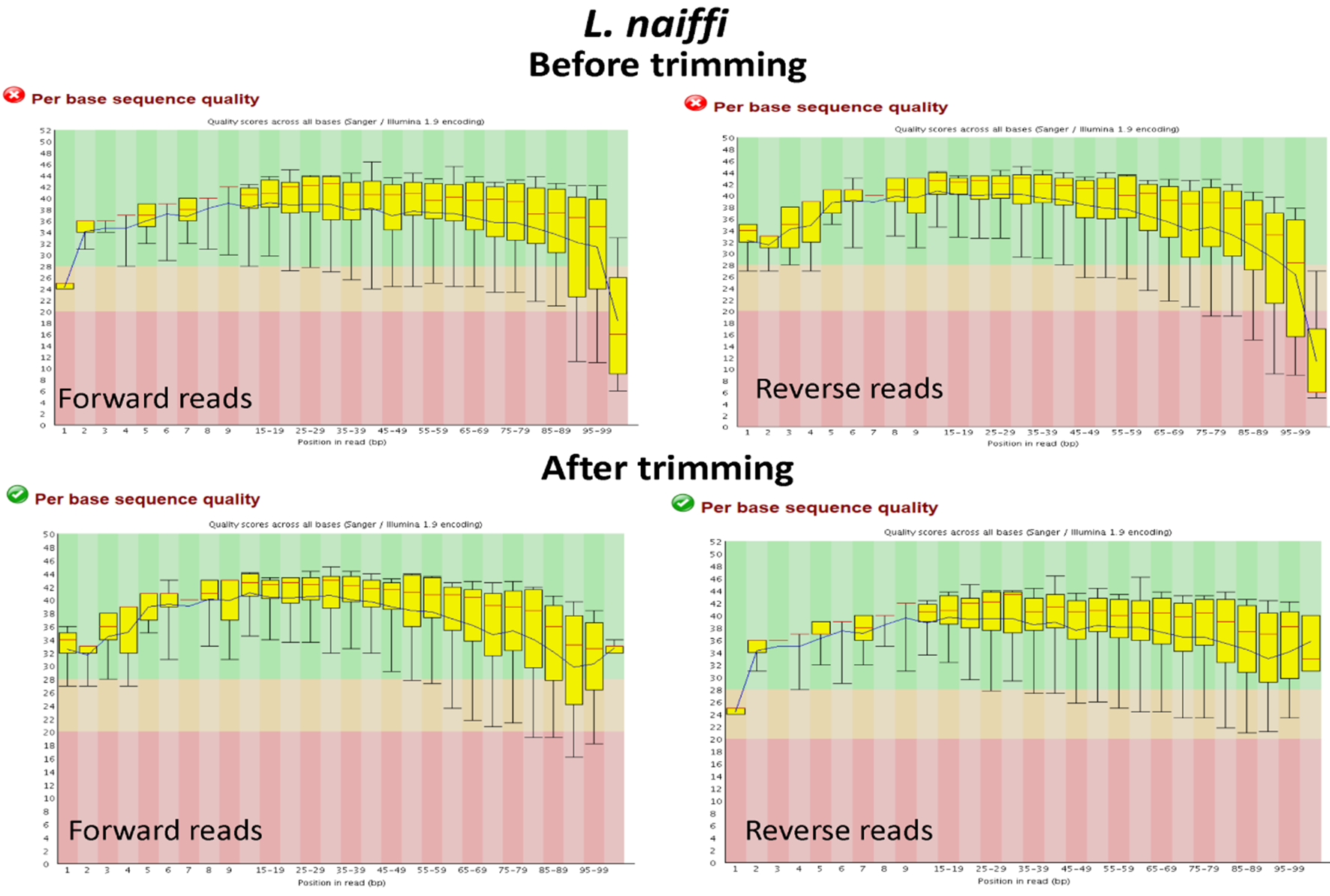

Figure S11: Per base sequence quality reported by FASTQC for *L. naiffi* LnCL223 reads before and after read trimming.

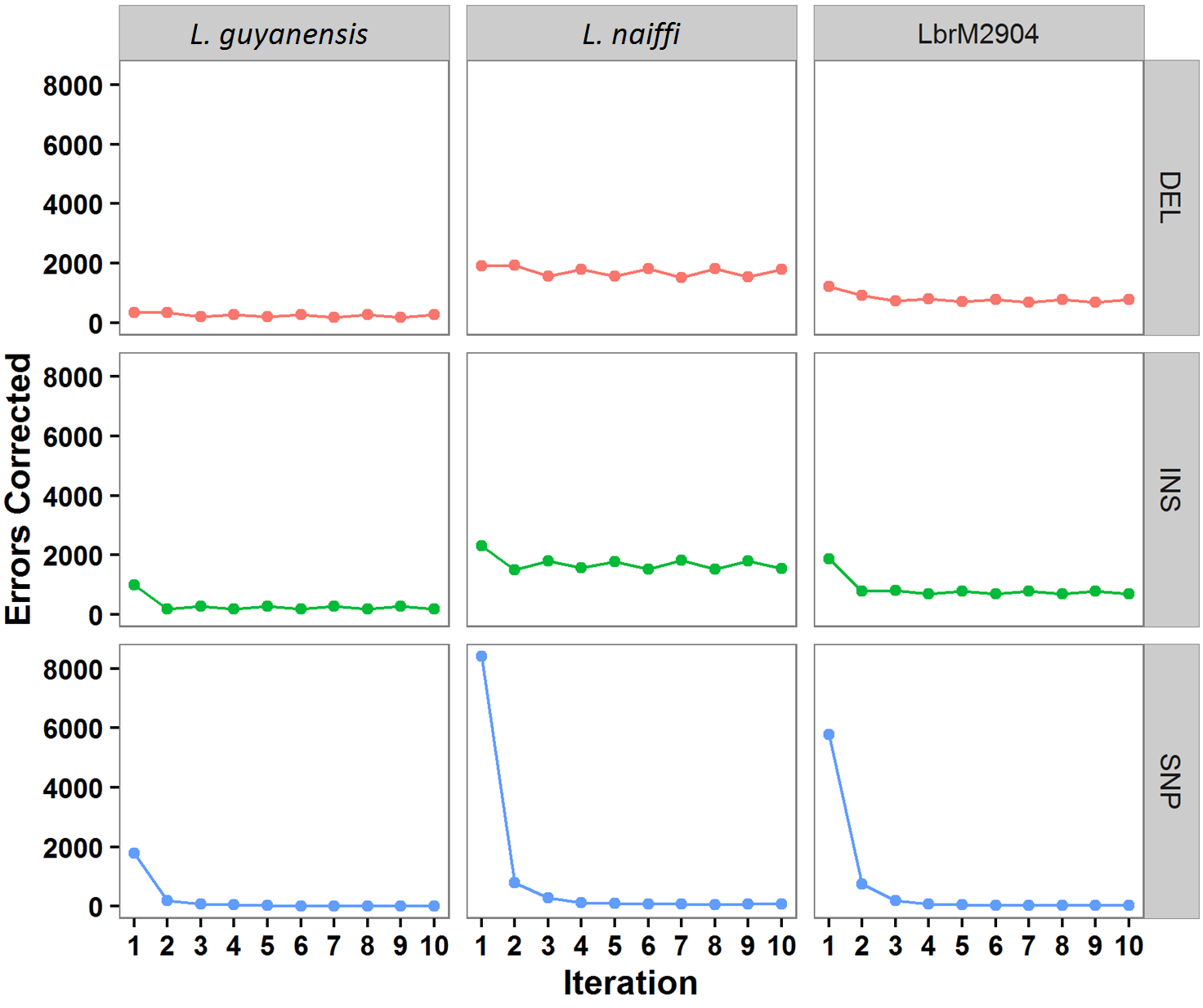

**Figure S12**: Number of deletions (DEL in red), short insertions (INS in green) and SNPs (blue) errors corrected at each iteration of ICORN for *L. guyanensis* LgCL085, *L. naiffi* LnCL223 and the control *L. braziliensis* genome (LbrM2904).

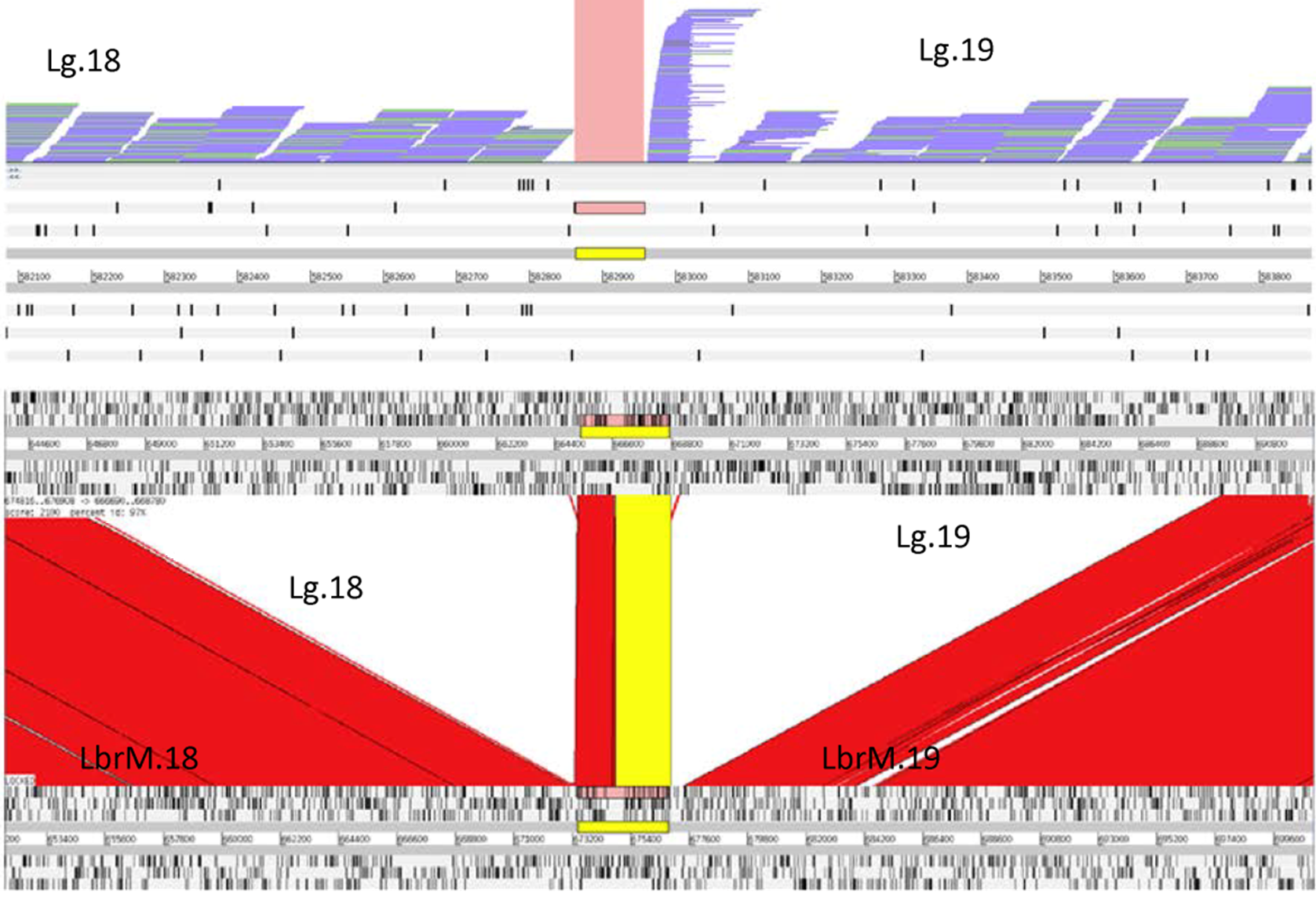

Figure S13: An example of the genome structure analyses. Top: *L. guyanensis* LgCL085 reads mapped to a concatenated version of chromosome 18 and 19 showed a gap bridged by no read pairs and a read pile-up at the start of chromosome 19. Bottom: Alignment of the same concatenated version of chromosome 18 and 19 with *L. braziliensis* M2904 chromosome 18 and 19. *L. guyanensis* LgCL085 chromosomes 18 and 19 were separated at this point.

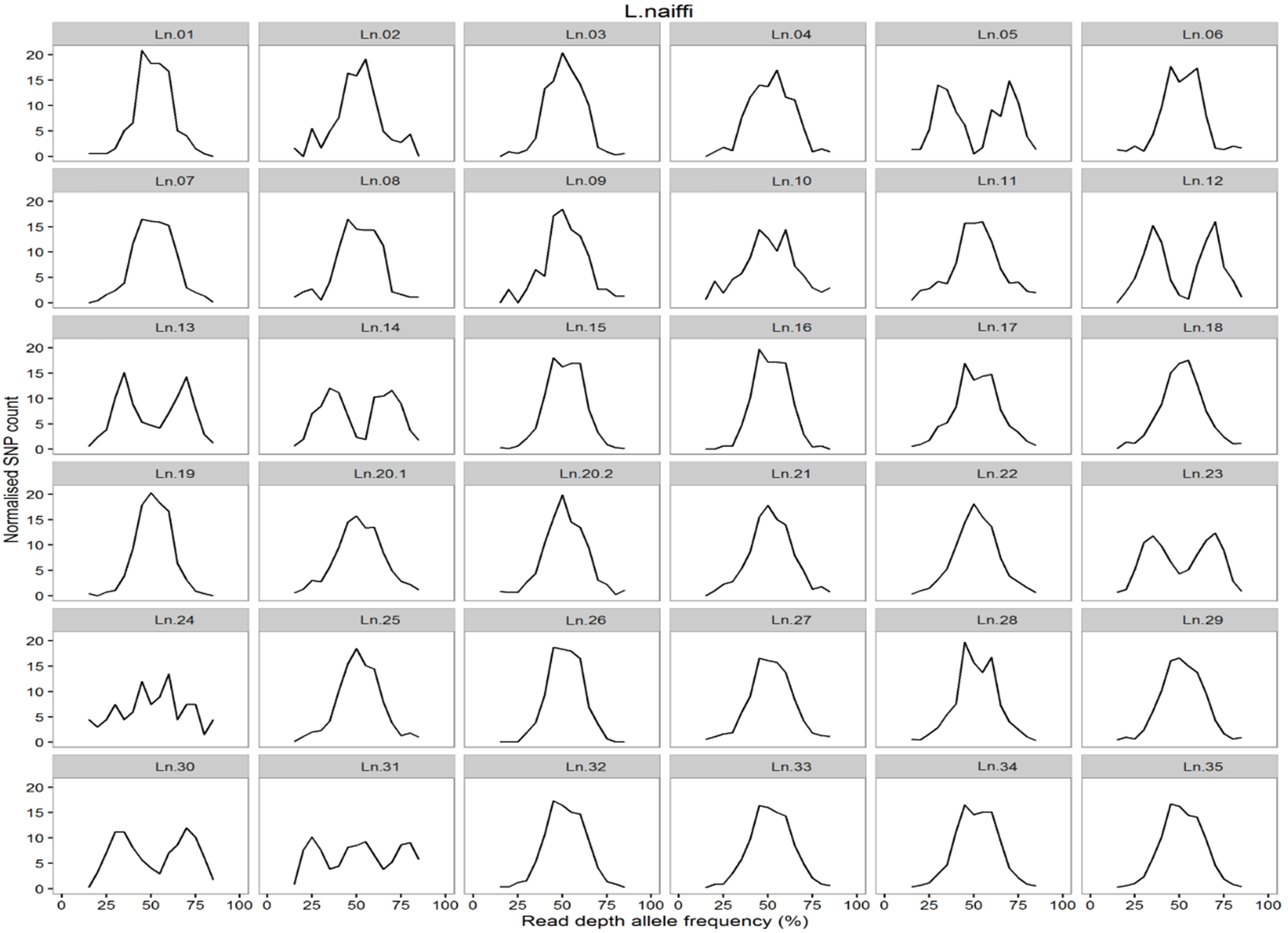

Figure S14: Read depth allele frequency distributions (RDAF) of each chromosome of *L. naiffi* LnCL223 determined using heterozygous SNPs called from reads mapped to its own assembly.

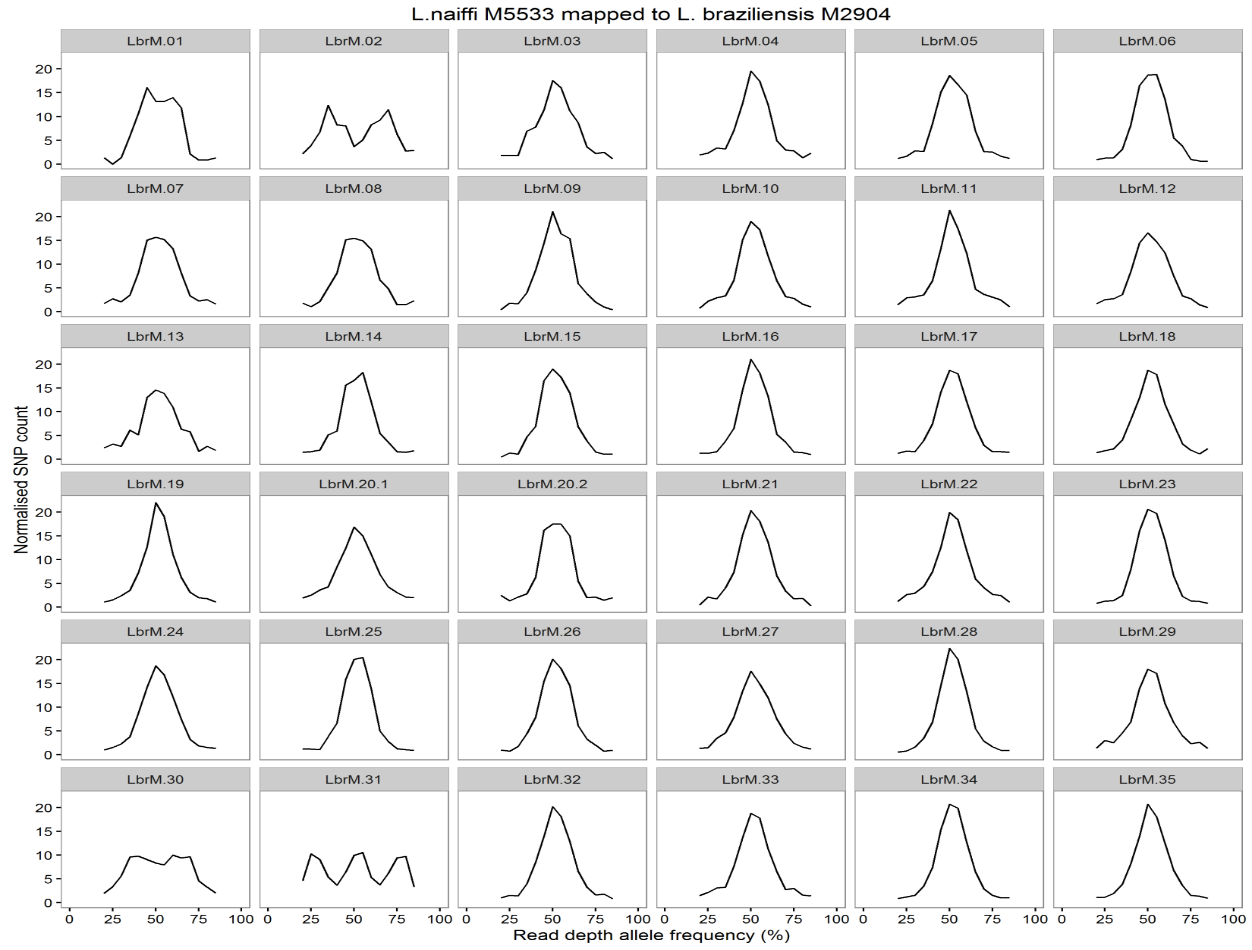

Figure S15: Read depth allele frequency distributions (RDAF) of each chromosome of *L. naiffi* M5533 determined using heterozygous SNPs from reads mapped to *L. braziliensis* M2904.

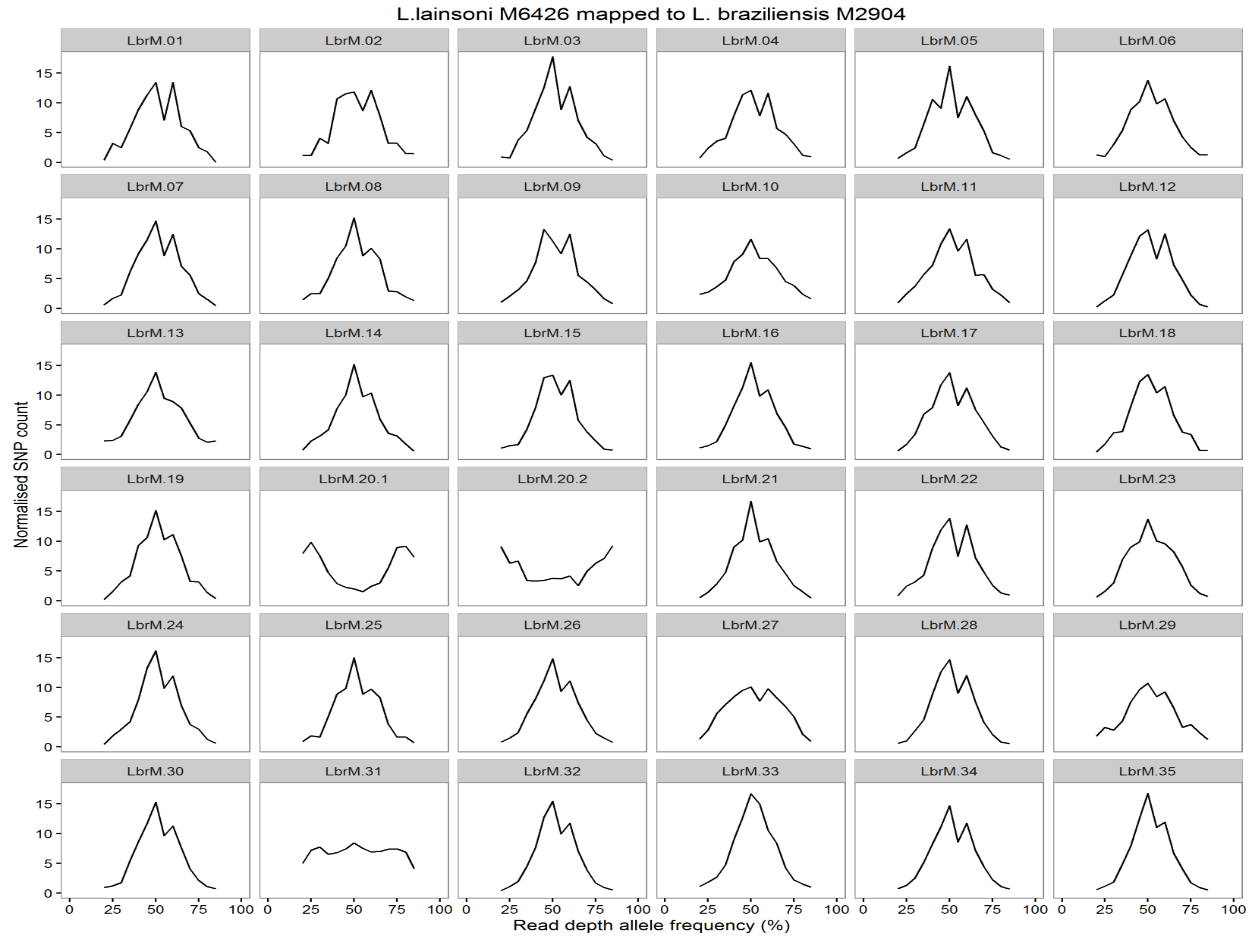

Figure S16: Read depth allele frequency distributions (RDAF) of each chromosome of *L. lainsoni* M6426 determined using heterozygous SNPs from reads mapped to *L. braziliensis* M2904.

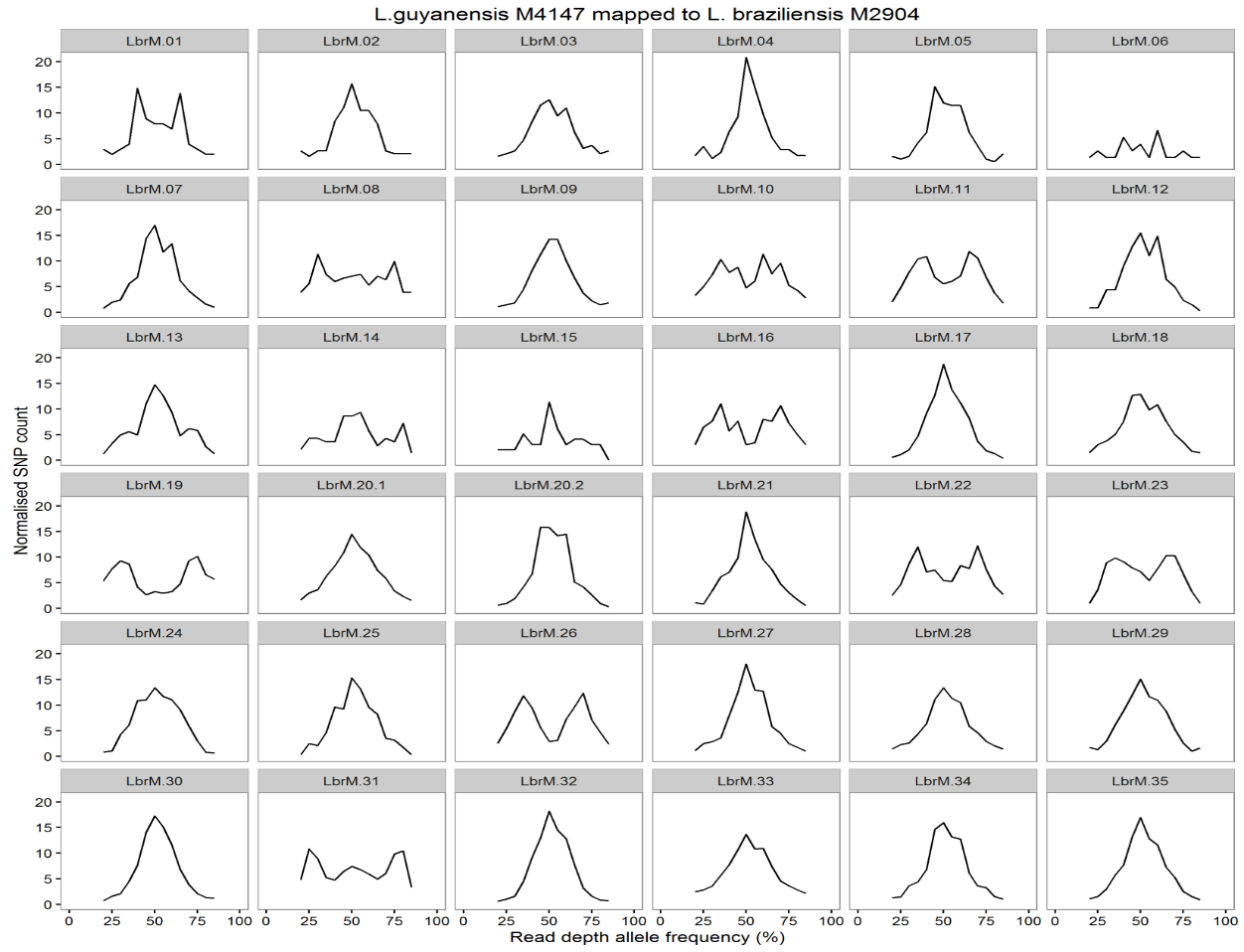

Figure S17: Read depth allele frequency distributions (RDAF) of each chromosome of *L. guyanensis* M4147 determined using heterozygous SNPs from reads mapped to *L. braziliensis* M2904.

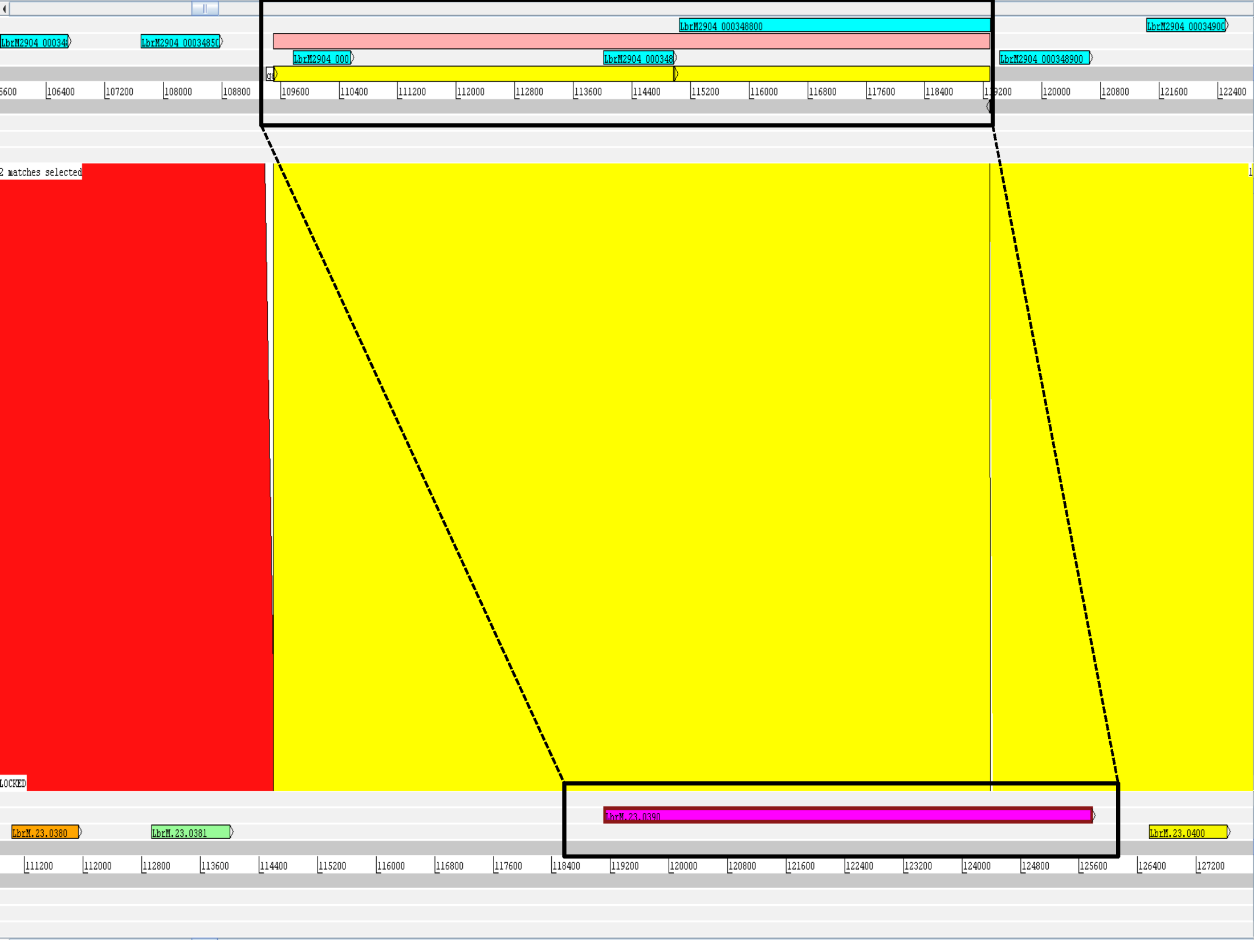

Figure S18: Incorrect models of the DCL1 gene on chromosome 23 of the *L. braziliensis* M2904 control genome (top) due to gaps. Only one gene model was present at the homologous locus on chromosome 23 of *L. braziliensis* M2904.

**Supplementary Methods**

***Viannia* comparative genome, annotation and proteome file sources**

The *L. braziliensis* M2904 reference genome and annotation (v3) EMBL files were from ftp://ftp.sanger.ac.uk/pub/project/pathogens/L_braziliensis/Archives/LbrM_v3_20110311/artemis/EMBL/Lbraziliensis/1/. The *L. panamensis* MHOM/PA/1994/PSC-1 genome and annotation files were download from NCBI accessions CP009370-404, the original 5,875,837 100 bp paired-end whole genome shotgun Illumina HiSeq 2000 reads from SRX681913, and its 7,748 protein coding sequences and GFF file from RefSeq ftp://ftp.ncbi.nlm.nih.gov/genomes/all/GCF_000755165.1_ASM75516v1/.

**kDNA assembly**

Sequences ≥ 1,000 bases in length that could not be assigned to chromosomes (bin sequences) were searched against BLAST databases of minicircle (753 sequences; “*kinetoplast AND minicircle AND leishmania*”) and maxicircle (152 sequences; query: “*kinetoplast AND maxicircle AND leishmania*”) sequences downloaded from Genbank using MegaBLAST. Hits were filtered to keep those with E-value < 0.01, bitscore > 100 and percentage identity > 40 to remove short hits. Sequences that had homology to both minicircle and maxicircle sequences were annotated in the bin sequences by adding ‘.kDNA.unassigned’ or ‘.kDNA.maxicircles’ or ‘.kDNA.minicircles’ to the ends of their headers.

**Manual correction of a false join of chromosomes 18 and 19**

The *L. guyanensis* LgCL085*,* *L. naiffi* LnCL223 and *L. braziliensis* control pseudo-chromosomes were visualized with the *L. braziliensis* M2904 chromosomes using the Artemis Comparison Tool (ACT). This manual verification process identified only one mistake in contiguity: a 101 Kb section at the 3’ end of chromosome 18 in *L. guyanensis* LgCL085, *L. naiffi* LnCL223 and the *L. braziliensis* M2904 control genome that was comprised of a part homologous to the start of chromosome 11 and a section similar to the start of chromosome 19 in the *L. braziliensis* reference. This was confirmed by alignment of the annotated genes using BLASTn and corresponded to a gap in coverage at the 3’ end of the reference chromosome 18 (Figure S13). On the basis that the *L. braziliensis* reference is the most accurate genome because it was created using long Sanger sequenced reads, each assembly was broken at this gap and joined to the end of chromosome 18 by a 100 bp gap. Likewise, the corresponding segment of chromosome 19 was transferred to the start of that chromosome with a 100 bp gap. The chunk of chromosome 11 was assigned as a bin contig because its 5’ end was not homologous with *L. braziliensis* M2904 chromosome 11. Gene model annotation and refinement at chromosome ends and variable regions remains an ongoing community challenge.

**Supplementary Results**

**Read-depth allele frequency (RDAF) distributions**

For *L. naiffi* LnCL223 chromosome 2 the RDAF distribution showed a large peak at 50% and two smaller peaks at 25% and 75% in comparison with that of chromosome 31, which had three approximately equal sized peaks (Figure S14). Six chromosomes (5, 12, 13, 14, 23 and 30) were trisomic and the remaining 27 were disomic. In *L. naiffi* M5533, only chromosome 2 was trisomic, as seen in both chromosome copy number and RDAF distribution plots (Figure 3, Figure S15), and all others were disomic, except for the tetrasomic chromosome 31. Chromosome 31 was tetrasomic in *L. shawi* M8408, *L. guyanensis* M4147, *L. guyanensis* LgCL085, *L. panamensis* WR120 and *L. lainsoni* M6426 (Figure 3). In *L. shawi* M8408, only chromosome 18 appeared trisomic (copy number of 2.8) based on coverage (Figure 3), but its RDAF profile indicated disomy (Figure S20). *L. panamensis* WR120 had six trisomic chromosomes (1, 5, 8, 23, 25 and 26) with all others disomic, bar tetrasomic chromosome 31 (Figure 3). *L. lainsoni* M6426 chromosome 33 was trisomic. Its chromosomes 20.1, 20.2, 23 and 27 exhibited values between disomic and trisomic states (2.5-3.0), but only chromosomes 20.1 and 20.2 showed RDAF peaks indicating trisomy, whereas 23 and 27 were disomic (Figure 3, Figure S16).

Chromosomes 1, 5, 6, 8, 13, 23, 26 and 35 of *L. guyanensis* LgCL085 were trisomic (eight trisomic chromosomes), chromosomes 7 and 31 were tetrasomic (copy numbers of 4 and 3.8 respectively) and the other 25 chromosomes were disomic (Figure 2). In *L. guyanensis* M4147, eight chromosomes were trisomic based on read depth analysis (Figure 3) and the RDAF distributions (Figure S17); these were chromosomes 8, 10, 11, 16, 19, 22, 23 and 26, so only chromosomes 8 and 26 were also trisomic in *L. guyanensis* LgCL085 (Figure 3). All other chromosomes in *L. guyanensis* M4147, with the exception of chromosome 31, were disomic.

**22 OGs exclusive to *Viannia* genomes**

The 22 OGs exclusive to *Viannia* genomes had many genes with no stated function. We identified those with annotated genes. Among these was one OG with beta tubulin and amastin-like surface protein genes with six genes in *L. naiffi* LnCL223, four in *L. braziliensis* and one in the other *Viannia* that has no orthologous genes in any other species outside *Leishmania*. The other three genes encoded a diacylglycerol kinase-like protein, a nucelobase transporter and an iron/zinc transporter protein-like protein that all had a haploid copy number of one. One encoded a eukaryotic translation initiation factor-like protein that had one copy in each genome and was otherwise only documented in *Trypanosoma brucei* and *Caenorhabditis elegans*. Another not shown was a possible additional copy of the ABCC7 protein PRP1 (pentamidine resistance protein 1, LbrM.31.1650 in *L. braziliensis*) gene that had one copy in each *Viannia* genome. The ABCC7 / PRP1 gene has multiple copies on *Viannia* chromosomes 23 and 31.

Among protease-related genes, only the M8 family metalloprotease leishmanolysin (GP63) array in OG5_126749 was amplified based on the 327 *L. naiffi* LnCL223, 334 *L. guyanensis* LgCL085 and 255 *L. braziliensis* M2904 control gene arrays (Table S23). OG5_129169 encoding a prenyl protease gene had elevated copy numbers in *L. braziliensis* and *L. naiffi*, likewise for a serine peptidase gene in OG5_126636 for *L. guyanensis* and *L. naiffi*, a cysteine peptidase gene in OG5_126607 for *L. braziliensis* and *L. guyanensis*, and a proteasome subunit gene in OG5_127562 in *L. naiffi*.

| **OG** | **Description** | **#Assembled genes in OG** | **Assembled genes in OG** | **OG haploid copy number** | **Species** |
| --- | --- | --- | --- | --- | --- |
| OG5_126749 | GP63, leishmanolysin, metallo-peptidase, Clan MA(M), Family M8 | 4 | LbrM2904_000106100, LbrM2904_000106200, LbrM2904_000106300, LbrM2904_000106700 | 5.31 | ***L. braziliensis control*** |
|  |  | 9 | LnCL223_100460, LnCL223_100470, LnCL223_100480, LnCL223_100490, LnCL223_100500, LnCL223_100510, LnCL223_100520, LnCL223_bin2370010, LnCL223_bin390010 | 55.74 | ***L. naiffi*** |
|  |  | 8 | LgCL085_100440, LgCL085_100450, LgCL085_100460, LgCL085_100480, LgCL085_100490, LgCL085_100500, LgCL085_bin1420010, LgCL085_bin1100020 | 32.7 | ***L. guyanensis*** |
| OG5_129169 | CAAX prenyl protease 2, peptidase with unknown catalytic mechanism (family U48) | 1 | LbrM2904_000436600 | 2.06 | ***L. braziliensis control*** |
|  |  | 1 | LnCL223_262610 | 2.26 | ***L. naiffi*** |
| OG5_126636 | ATP-dependent Clp protease subunit, heat shock protein 100 / 78 (HSP100/HSP78), serine peptidase | 2 | LnCL223_272730, LnCL223_291340 | 4.27 | ***L. naiffi*** |
|  |  | 2 | LgCL085_272680, LgCL085_291340 | 3.24 | ***L. guyanensis*** |
| OG5_126607 | cysteine peptidase A (CPA/CBA), cathepsin L-like protease | 2 | LbrM2904_000793200, LbrM2904_000248500 | 3.17 | ***L. braziliensis control*** |
|  |  | 2 | LgCL085_191470, LgCL085_080840 | 4.03 | ***L. guyanensis*** |
| OG5_127562 | 20s proteasome beta 7 subunit | 1 | LnCL223_2013970 | 2.55 | ***L. naiffi*** |

**Table S23**. Gene arrays in the *L. braziliensis* control, *L. guyanensis* LgCL085 and *L. naiffi* LnCL223 for gene related to proteases with haploid copy numbers of two or more per OG.

**Gene annotation version and tools**

A large number of bin contigs were annotated as kDNA minicircles: 24 for *L. guyanensis* LgCL085 (total length 961,641 bp) and four as unassigned kDNA (total length 5,090 bp); 23 for *L. naiffi* LnCL223 (98,031 bp); and ten for the *L. braziliensis* M2904 control assembly (158,544 bp). No bin contigs had homology to maxicircles.

The genes in OGs annotated on the *L. braziliensis* control M2904 by Companion but not on the original published *L. braziliensis* M2904 annotation are in Table S19.

A gene encoding a methyltransferase-domain containing protein (LnCL223_2021430 in OG5_129552) in *L. naiffi* LnCL223 may be a valid protein coding gene because it was in the most recent unpublished *L. braziliensis* genome annotation and had orthologs in other eukaryotes, but it was absent in the published version (Table S13).

Two genes previously exclusive to *L. adleri* (Table S20) were in *L. naiffi* LnCL223, *L. guyanensis* LgCL085 and *L. braziliensis*, perhaps due to differing annotation protocols between studies - the most powerful tool called Companion was used, unlike previous work.

The amplifications on *L. guyanensis* LgCL085 and *L. naiffi* LnCL223 of loci at least 10 Kb provided details on the spectrum of CNVs in *Viannia* genomes (Table S21).

***L. guyanensis* LgCL085 and *L. naiffi* LnCL223 RNAi pathway genes**

RNA interference (RNAi) is a post-transcriptional gene silencing mechanism initiated by short double-stranded RNA (dsRNA). RNAi is present in *Trypanosoma brucei* [1,2], *L. braziliensis* [3,4]*, L. guyanensis* LgCL085, *L. panamensis* and the non-parasitic species *Crithidia fasciculata* [3], but not in the *Leishmania* or *Sauroleishmania* subgenera. In this pathway, dsRNA is converted to small interfering RNA (siRNA) by Dicer. Five RNAi genes were in *L. braziliensis*: cytoplasmic Dicer like 1 (DCL1, LbrM.23.0390)*,* nuclear Dicer like 2 (DCL2, LbrM.25.1020) [1], Argonaute 1 (AGO1, LbrM.11.0360) [4] and RNA Interference Factors 4 and 5 (RIF4 and RIF5, LbrM.35.6220 and LbrM.33.0190) [5]*.* Each of these genes had orthologs in *L. guyanensis* LgCL085*, L. naiffi* LnCL223, the *L. brazilensis* control assembly, *L. panamensis* PSC-1*, L. peruviana* PAB-4377 and *L. peruviana* LEM1537 (Table S22) and (as a negative control) none in *L.* *adleri* HO174 or *L. tarentolae* as expected. A hypothetical gene (LbrM.29.0560) associated with siRNA production (GO: 0030422) was also in an orthologous group with the AGO1 gene. The RIF5 gene had orthologs in *L. major,* *L. infantum*, *L. mexicana* and *L. donovani*. AGO1 had mutated orthologs in *L. major* (LmjF.11.0570) and *L. infantum* (LinJ.11.0500). *L. guyanensis* LgCL085 and *L. naiffi* LnCL223 orthologs with complete ORFs were present at the homologous regions of *L. braziliensis* and had one haploid copy of each gene. The *L. braziliensis* M2904 control genome had three copies of DCL1 in contrast with one copy in the published assembly. This was caused by a single ‘N’ base and an 11 bp gap before the 5’ end of the gene that resulted in three separate annotated genes instead of just one (Figure S18).
